# Transcriptomic Insights into the Epigenetic Modulation of Turnip Mosaic Virus Evolution in *Arabidopsis thaliana*

**DOI:** 10.1101/2024.06.18.599562

**Authors:** María J. Olmo-Uceda, Silvia Ambrós, Régis L. Corrêa, Santiago F. Elena

## Abstract

Plant-virus interaction models propose that a virus’s ability to infect a host genotype depends on the compatibility between virulence and resistance genes. Recently, we conducted an evolution experiment in which lineages of turnip mosaic virus (TuMV) were passaged in *Arabidopsis thaliana* genotypes carrying mutations in components of the DNA methylation and the histone demethylation epigenetic pathways. All evolved lineages increased infectivity, virulence and viral load in a host genotype-dependent manner. To better understand the underlying reasons for these evolved relationships, we delved into the transcriptomic responses of mutant and WT plant genotypes in mock conditions and infected with either the ancestral or evolved viruses. Such a comparison allowed us to classify every gene into nine basic expression profiles. Regarding the targets of viral adaptation, our analyses allowed the identification of common viral targets as well as host genotype-specific genes and categories of biological processes. As expected, immune response-related genes were found to be altered upon infection. However, we also noticed the pervasive over-representation of other functional groups, suggesting that viral adaptation was not solely driven by the level of expression of plant resistance genes. In addition, a significant association between the presence of transposable elements within or upstream the differentially expressed genes was observed. Finally, integration of transcriptomic data into a virus-host protein-protein interaction network highlighted the most impactful interactions. These findings shed extra light on the complex dynamics between plants and viruses, indicating that viral infectivity depends on various factors beyond just the plant’s resistance genes.

## Introduction

Plants and viruses engage in intricate interactions that trigger defense and counter-defense mechanisms, often resulting in coevolution and reciprocal adaptation. Plant immune processes play a pivotal role in antiviral defense (Soosaar, Burch-Smith and Dinesh-Kumar 2005; Zhou and Zhang 2020). These antiviral factors can be classified into two groups: basal, which are preexisting and restrict viral propagation within and between cells, and inducible, which are activated upon infection and hinder systemic movement and replication. Inducible mechanisms involve genes that induce broad-scale changes in plant physiology through various signaling pathways, while basal mechanisms correspond to alleles of cell proteins whose interaction with viral factors is altered (Carr, Lewsey and Palukaitis 2010). These changes include local cell apoptosis (Loebenstein 2009), upregulation of nonspecific responses against pathogens throughout the entire plant (systemic acquired resistance -SAR- and induced systemic resistance -ISR-) (Kachroo, Chandra-Shekara and Klessig 2006; Carr, Lewsey and Palukaitis 2010), and the activation of RNA-silencing-based resistance, which plays a role in both basal and inducible mechanisms (Voinnet 2001; Carr, Lewsey and Palukaitis 2010; López-Gomollon and Baulcombe 2022).

RNA-based immunity is initiated by recognizing and degrading double-stranded RNAs (dsRNA). During viral replication, Dicer-like (DCL) proteins break down viral dsRNAs into small RNAs (sRNA), which are then loaded into Argonaute (AGO) proteins to silence complementary RNAs (Voinnet 2001). *Arabidopsis thaliana* genome has four DCL genes, with the majority of viral siRNAs depending on the DCL4 protein. However, strong antiviral defense may also need the hierarchical activity of DCL2 and DCL3 (García-Ruiz et al. 2010). Enhancing the silencing response involves RNA-dependent RNA polymerases (RDR), which produce additional dsRNAs from the target (Borges and Martienssen 2015).

These RNA-mediated defense mechanisms are part of a larger and evolutionarily conserved system that regulates gene expression and manages transposable elements (TE) through epigenetic modifications of DNA or histones (Hung and Slotkin 2021). DNA methylation, observed in all sequence contexts (CG, CHG, and CHH; with H being any nucleotide except G), is initiated by a well-studied mechanism known as RNA-directed DNA methylation (RdDM). This process utilizes sRNAs produced from the borders of TEs to restrict their expression and influence neighboring gene expression (Liu et al. 2022). RdDM relies on several components, including DCL3, METHYLTRANSFERASE DOMAINS REARRANGED METHYLASE 2 (DRM2), RNA polymerase IV (POLIV), and RNA polymerase V (POLV) (Böhmdorfer et al. 2016). Various non-canonical pathways also contribute to the canonical RdDM pathway, such as DCL-independent mechanisms, POLII-derived mRNAs, or RDR6-derived dsRNAs (Cuerda-Gil and Slotkin 2016). Epigenetic marks are also copied during replication by maintenance DNA methyltransferases like the plant-specific CHROMOMETHYLASE 3, and they require chromatin remodelers such as DECREASED DNA METHYLATION 1 (DDM1) to control gene expression and TE mobilization in sRNA-independent mechanisms (Bond and Baulcombe 2014). Histone modification is also linked to DNA methylation marks. For instance, histone H3 trimethylation of lysine 9 (H3K9m3) is associated with TE repression, while H3K4m3 stimulates gene expression. Both DNA methylation and histone modifications are reversible, with proteins like REPRESSOR OF SILENCING 1 (ROS1), INCREASE IN BONSAI METHYLATION 1 (IBM1), and JUMONJI14 (JMJ14) involved in demethylation of DNA, H3K9, and H3K4, respectively (Gong et al. 2002; Saze et al. 2008; Lu et al. 2010). Under stressful conditions or in epigenetically-deficient mutants, the chromatin environment around TEs and genes can be altered, affecting their expression (Lloyd and Lister 2021).

The link between the loss of DNA methylation factors and resistance or susceptibility to RNA virus infection has received limited attention. Research using turnip crinkle virus (Diezma-Navas et al. 2019), two tobamoviruses (Leone et al. 2020), and turnip mosaic virus (TuMV) (Corrêa et al. 2020) has revealed that *A. thaliana* mutants for various RdDM factors, chromatin remodelers, and histone modifiers exhibited varying susceptibility to infections based on the presence of repressive marks. Additionally, evolution experiments involving mutants for innate immunity pathways and basal and inducible defense pathways have shown that viral populations rapidly adapt to the novel selective pressures imposed by mutations in RdDM genes (Navarro et al. 2022; Ambrós et al. 2024) and histone modification genes (Ambrós et al. 2024), indicating a direct role of these genes in the infection cycle of RNA viruses.

A consistent observation in evolution experiments with potyviruses is that the transcriptomic response of *A. thaliana* changes as viral lineages became better adapted to the new host (Agudelo-Romero et al. 2008; Hillung et al. 2016; Cervera et al. 2018; Corrêa et al. 2020). Indeed, differences in transcriptomic responses arise at different levels of *A. thaliana* genetic diversity, among ecotypes but also among mutant genotypes. To investigate the role of epigenetic regulation of host genes involved in defense responses in the probability of an RNA virus establishing successful infections after spilling-over from a reservoir to a naïve host, Ambrós et al. (2024) evolved TuMV lineages in *A. thaliana* plants with mutations in DNA methylation and histone demethylation pathways. They found that evolved viral lineages become more infectious, virulent, and reached higher viral loads. Interestingly, the extent of these phenotypic changes depended on the mutated epigenetic pathway. Indeed, *jmj14* with affected histone methylation selected for more virulent viruses, while mutants unable of producing siRNAs (*dcl2 dcl3 dcl4*) and thus particularly sensitive to infection, selected for less virulent viral strains. We hypothesize that given the expected large-scale effects of mutations in epigenetic regulatory pathways in plant defenses and cellular homeostasis, virus evolution and adaptation to the novel host will be affected. This impact can be either direct, if epigenetic factors directly interact with virus factors, or indirect if epigenetic regulation modifies the expression of plant resistance genes that ultimately interact with viral factors or induce a strong physiological dysregulation that represents a different cellular environment for the virus. Here, to further characterize the role of DNA methylation and histone modification pathways in the adaptation of TuMV, we sought differences in whole-genome transcriptomic responses to infection with the ancestral TuMV and derived viral lineages in their corresponding local host genotypes (fig. 1).

**Figure 1.**
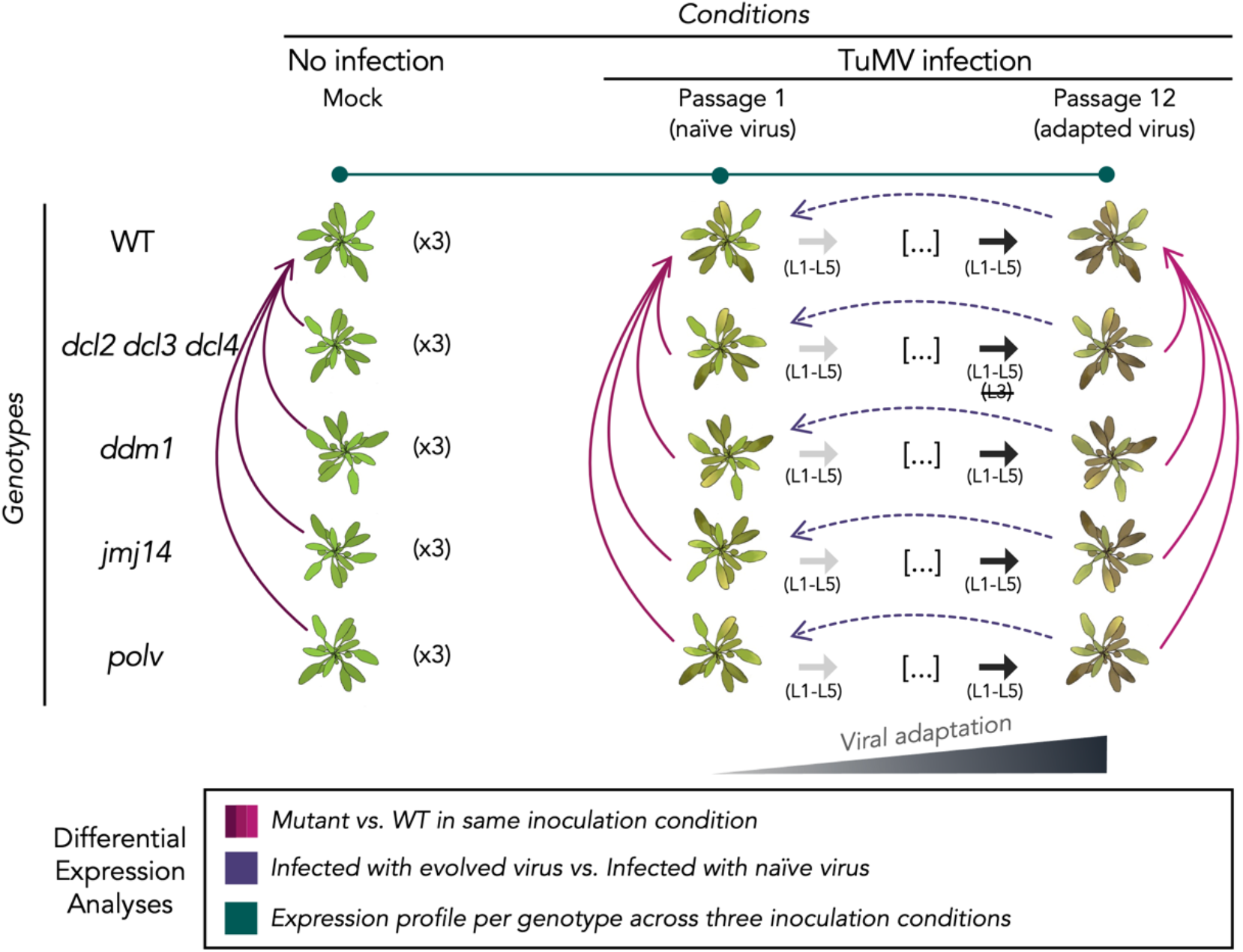
Schematic representation of the evolution experiment and the three different contrasts performed on the transcriptomic data. Considering three inoculation conditions: mock (no infection), infection with the ancestral virus (P1) and infection with the evolved viral lineages (P12) and the five different plant genotypes, three blocks of differential expression analyses were performed (represented by arrows, with the arrowhead pointing to the baseline of the contrast). Bourgogne: differences between each mutant and the WT in each condition. Purple: differences between the infection with adapted and the naïve virus for each genotype. The lineages were used as block variables. Green: the differential expression analyses between infection with naïve and mock and the differences between adapted and naïve infection (purple lines) led us to classify each gene expression in nine different profiles.

## Results and Discussion

Fig. 1 illustrates the design of the evolution experiment, with indication of the specific contrasts made across transcriptomic profiles. RNA-Seq was conducted for plants of four mutants plus wild-type (WT) infected with the corresponding viral lineages isolated after passages 1 (P1), hereafter referred to as the ancestral stage, and 12 (P12), henceforth as the evolved lineages. In addition, the transcriptomes of mock-inoculated plants from the five genotypes were also determined. The *dcl2 dcl3 dcl4* triple mutant exhibits dysregulation of many RNA silencing mechanisms, such as the main bulk of antiviral siRNAs, RdDM, and endogenous siRNAs derived from dsRNA produced by RDR6. This mutant has been shown to be highly susceptible to virus infection (García-Ruiz et al. 2010). Both *ddm1* and *polv* mutants have altered DNA methylation-related mechanisms. DDM1 is a master regulator of TEs, and protein deficiencies cause a steady loss of RdDM-independent DNA methylation and large transposition events (Zemach et al. 2013). POLV plays a critical role in the process of RdDM. Its transcripts are responsible for initiating CHH methylation on both strands of DNA, specifically targeting short TEs and the edges of long ones (Böhmdorfer et al. 2016). In *polv* mutants, a significant decrease in non-CG methylation across the entire genome is observed, accompanied by the disruption of various stress-related genes (Stroud et al. 2013; Corrêa et al. 2024). The *jmj14* mutant has been shown to have higher levels of H3K4 activation marks in its direct targets (Greenberg et al. 2013).

### Summary of differentially expressed genes amongst infection treatments and plant genotypes

First, we analyzed the differences in the transcriptomes of the mutants with respect to the WT in each inoculation condition (fig. 2 and supplementary fig. S1, Supplementary Material online). These contrasts represent a first approach to discover differences in the way evolution has changed the interaction of evolved viruses with their host genotypes deficient in epigenetic components relative to their interaction with WT plants that are fully competent in all epigenetic regulation pathways. Fig. 2A shows the distribution of the number of differentially expressed genes (DEGs) for each of the four mutant plant genotypes compared to WT plants, non-inoculated. This distribution sets the baseline for the overall effect of the different mutations in plant transcriptomes in absence of infection. The plant genotype showing the largest amount of genotype-specific DEGs was *dcl2 dcl3 dcl4* (2073), followed by *ddm1* (809), *polv* (462), and *jmj14* (258) (supplementary fig. S1B, Supplementary Material online). Indeed, the number of genes altered in common by the three mutants affecting DNA methylation-related pathways was 313, 21.32% larger than the number of genes altered by the histone modification mutant. One-hundred sixty DEGs were in common for the four infected mutants. Remarkably, these genes were grouped into two large, well-balanced, clusters of over- and under-expressed genes highly consistent across the four mutant plants (fig. 2A) with only two genes presenting a different expression pattern between the mutants (left panel in supplementary fig. S1A, Supplementary Material online).

**Figure 2.**
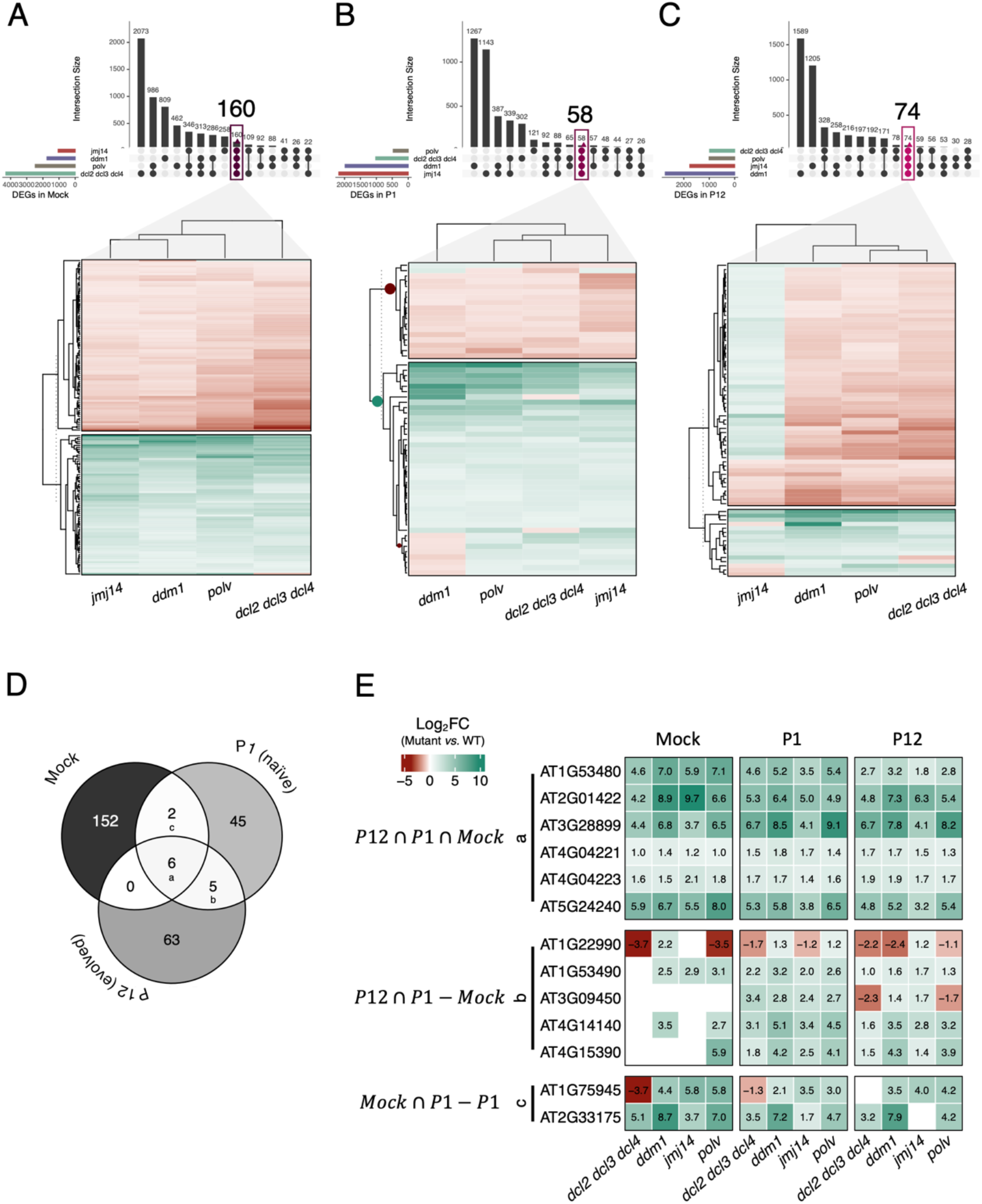
Distribution of differentially expressed genes (DEGs) compared to WT per plant genotype for mock-inoculated plants (A), plants inoculated with the ancestral virus (B) and plants inoculated with the evolved lineages (C). Expression clustering for DEGs shared by all mutant plants are indicated in each case. (D) Distribution of DEGs shared by all the mutants (core genes) in mock-inoculated and plants infected with the ancestral (P1) and evolved (P12) viruses. (E) Gene expression (log_2_*FC* represent mutant *vs.* WT expression by inoculation condition) of the three overlapping sets of core genes in (D), only significant DEGs (adjusted *P* < 0.05) with |log_2_*F*C| > 1 expression levels are shown, rest in white.

Second, we sought to evaluate the differences with WT in the plants infected with the ancestral P1 viral populations (fig. 2B). The distribution of mutant-specific DEGs has significantly changed relative to the one observed for mock-inoculated plants (χ^2^ = 3828.937, 14 d.f., *P* < 0.001): now *ddm1* plants show the largest number of DEGs (1267), followed by *jmj14* (1143), *dcl2 dcl3 dcl4* (302), and *polv* (121). The subset of DEGs in common for the three infected DNA methylation-related mutants was now 27, which represents 97.64% less than the number of DEGs affected by the histone modification infected *jmj14* mutant. A total of 58 DEGs were shared by all four mutant plants after infection. These DEGs group in two major clusters, with an excess of overexpressed genes, although *ddm1* plants showed a group of eight underexpressed DEGs that were overexpressed in all the other mutants (fig. 2B, intermediate panel in supplementary fig. S1A, Supplementary Material online).

Third, we compared the number of DEGs in plants infected with the evolved lineages (fig. 2C). In this case, the relevant comparison is between the number of DEGs observed for each plant genotype and ancestral and evolved viruses. Significant differences have been generated during the course of evolution (χ^2^ = 733.467, 14 d.f., *P* < 0.001), mostly driven by an increase in the relative number of *ddm1* (percentage deviation 4.9%) and *polv* (20.8%) specific DEGs along with a reduction in the relative numbers of *jmj14* (3.3%) and *dcl2 dcl3 dcl4* (26.7%). In this case, 74 DEGs were shared by all four infected mutant plant genotypes. These DEGs are clustered into two main categories: 17 over- and 57 under-expressed. However, the behavior of infected *jmj14* plants was remarkably different from the other mutants, with most of these DEGs being overexpressed (fig. 2C, right panel in supplementary fig. S1A, Supplementary Material online). Overall, the percentage of DEGs shared by all mutants incresed from the 1% in mock, to 21 % in plants infected with the ancestral viruses and to 72% in plants inoculated with the evolved ones (fig. S1B, Supplementary Material online), being the difference highly significant (χ^2^ = 109.360, 2 d.f., *P* < 0.001).

Finally, we evaluate the number of mutant-specific responses integrated with the degree of transcriptome perturbation, observing a differential pattern among mutants (supplementary fig. S1B, Supplementary Material online). The direction of change, however, differed among genotypes: (*i*) the level of perturbation is higher in mock conditions than in infection for *dcl2 dcl3 dcl4* and *polv*, the most permissive genotypes (Ambrós et al., 2024) but not in the rest. (*ii*) The percentage of mutant-specific DEGs was higher in *ddm1* and *jmj14* mutants during infection (above 50%). And (*iii*) the percentage of mutant-specific DEGs seems to stabilize in *ddm1* (from 46% of specificity in mock conditions to 64% and 58% in evolved viral lineages) while continues growing in *jmj14* (from 24% in mock to 53% in ancestral and 68% in evolved viruses). In agreement with the observation that TuMV rate of evolution was faster in *jmj14* (Ambrós et al. 2024), these results confirm the stronger selective pressure that this mutant imposes on the virus.

For the sake of simplicity, hereafter we will focus on a reduced core of DEGs shared by all plant genotypes: six among all three treatments, five between infected plants but not with mock-inoculated plants and two that are shared between mock-inoculated and ancestral-inoculated plants (fig. 2D-E).

### Epigenetic mutants exhibit changes in a reduced set of common genes

Fig. 2D illustrates the overlapping among the 13 core DEGs. The six overexpressed DEGs shared by all plant genotypes, regardless of their infection status, are shown in fig. 2E(a). These six loci (supplementary table S1, Supplementary Material online) can be seen as nonspecific responses of plants, regardless of their epigenetic context, to the inoculation process and growth conditions. This list includes *METHIONINE OVERACCUMULATION 1 (MTO1) RESPONDING DOWN 1* (*MRD1*), a gene that encodes for a protein involved in down-regulation of methionine biosynthesis, and is essential for salicylic acid (SA)-mediated defense (Singh et al. 2022), a ribosomal protein L34e superfamily member (AT3G28899), the ubiquitin family protein gene *1-PHOSPHATIDYLINOSITOL 4-KINASE C3* (*PI4KC3*), involved in regulation of flower development and responses to abscisic acid (Akhter et al. 2015) and three noncoding RNAs (AT2G01422, AT4G04221 and AT4G04223). Interestingly, *MRD1*, *PI4KC3*, AT2G01422, AT3G28899, and AT4G04223 have previously been shown to be directly regulated by RdDM (Kurihara et al. 2008; Blevins et al. 2014; Corrêa et al. 2024).

Fig. 2E(b) shows the expression pattern of the five core DEGs shared only by infected plants irrespective of virus’ evolution status (supplementary table S1, Supplementary Material online). These five DEGs would represent shared responses to infection. This list includes genes related to stresses (AT1G22990, AT3G09450 and AT4G15390), meiosis (AT1G53490) and the methyltransferase *MET2*. AT1G53490, AT4G15390 and *MET2* are consistently induced upon infection with both the ancestral and evolved viruses in all plant genotypes. Changes of expression direction due to virus evolution is observed for the gene AT1G22990 in *ddm1*, *jmj14* and *polv*, indicating a potential role in the adaptation process in these particular genotypes. The same holds true for the gene AT3G09450 in the DNA methylation-related *dcl2 dcl3 dcl4* and *polv* mutants. Among the five genes, one (AT1G53490) is predicted to be directly regulated by POLV (Corrêa et al. 2024).

Two core DEGs are shared by mock- and ancestral-inoculated plants, but not with the evolved-inoculated ones (fig. 2D; supplementary table S1, Supplementary Material online). Therefore, these two DEGs can be seen as either targets or drivers of TuMV adaptation to specific mutant plant genotypes. AT1G75945 and AT2G33175 both encode for mitochondrial membrane-associated hypothetical proteins that both were overexpressed compared to WT plants, except in *dcl2 dcl3 dcl4* plants, in which the first one was underexpressed (fig. 2E(c)). Notably, the ∼4-fold increase in expression observed in *polv* plants agrees with the previous observation by Corrêa et al. (2024).

### Impact of viral adaptation into plant responses

Once the differences between mutants and WT plants in their response to infection where explored, we focused on the expression changes elicited by the adaptation of the viral lineages to their corresponding local host genotypes [fig. 1 purple lines: infected with evolved viral lineages (P12) *vs.* infected with their corresponding ancestors (P1)]. In this context, by up-regulated we meant significantly more expressed in plants infected with the evolved lineages with respect to plants infected with their corresponding ancestors; by down-regulated we meant the opposite. To simplify wording, we will refer to these genes as activated or repressed by the viral adaptation. The total number of virus-adaptation DEGs was larger in *jmj14* and *polv* than in WT (fig. 3A) while, at the other extreme, *dcl2 dcl3 dcl4* and *ddm1* showed a similar degree of disruption than WT plants (fig. 3A). However, it is noticeable that in all four mutants, the distributions of differentially activated and repressed genes were different than observed in WT plants (Fisher’s exact tests, *P* < 0.001). Overall, in all mutant plant genotypes, the number of repressed genes was larger than in WT plants. The situation was more variable for activated genes, with *dcl2 dcl3 dcl4*, *ddm1* and *polv* showing less than WT plants but *jmj14* showing more.

**Figure 3.**
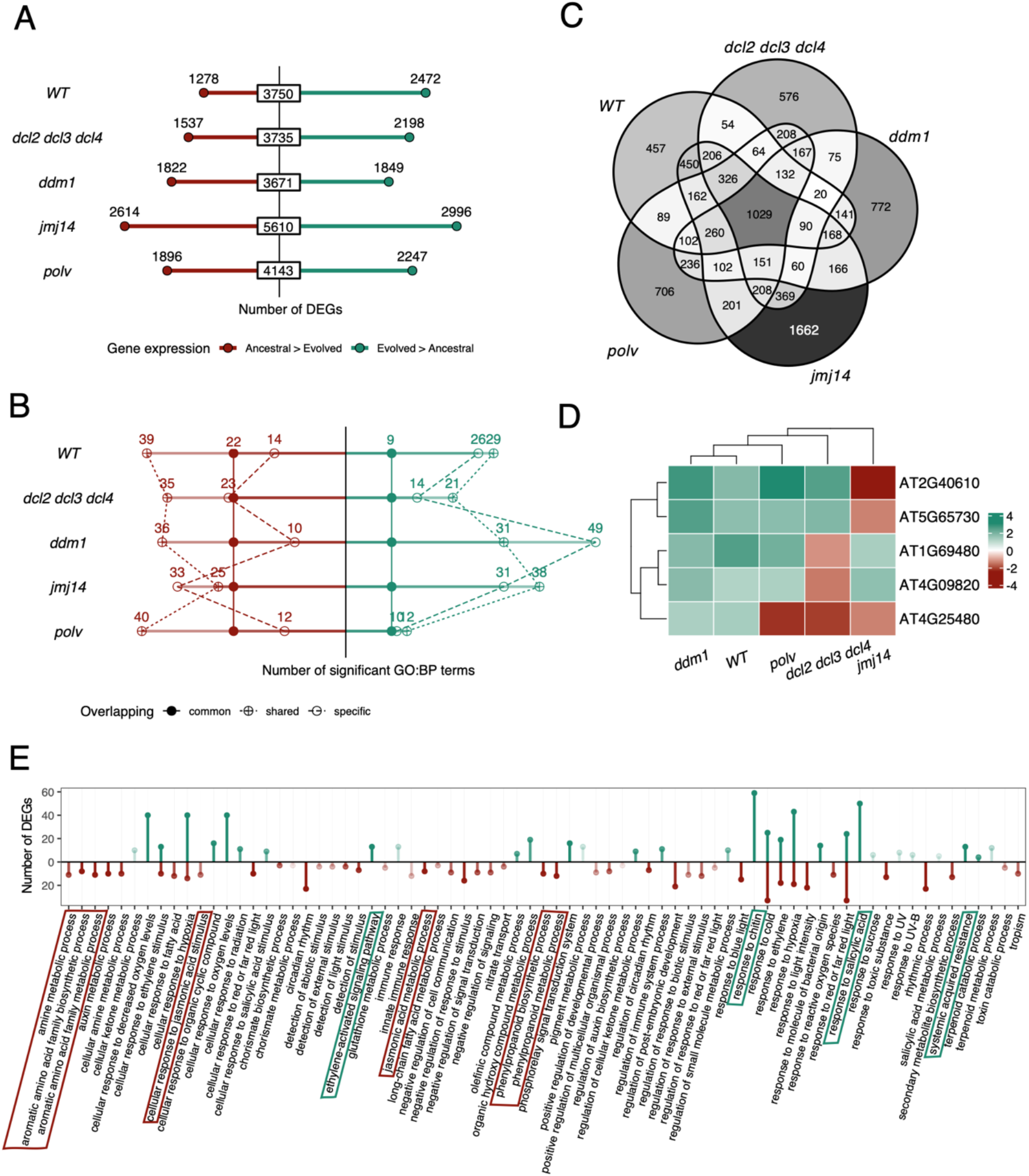
Characterization of DEGs between plants infected with the ancestral and evolved viruses, per plant genotype. (A) Number of repressed (red) and activated (green) DEGs elicited by viral adaptation in each plant genotype. (B) Number of GO biological processes terms within the repressed (red) and activated (green) DEGs enriched per plant genotype, indicating the number of common to all plant genotypes (solid symbols), shared by more than one plant genotype (crossed symbols), and specific of a single plant genotype (open symbols). (C) Distribution of DEGs across plant genotypes. (D) Only five of the 1029 DEGs common to all genotypes, present an opposite gene expression. The color scale represents the level of expression in log_2_*FC* as in (A) and (B). (E) GO biological processes terms from DEGS in common indicated in panel (C). In green, enriched categories within the activated DEGs, in red enriched categories within the repressed DEGs (adjusted *P* < 0.05).

A functional enrichment analysis (GO: biological process) of the DEGs separated by its regulation revealed some categories common to all the genotypes (supplementary fig. S2, Supplementary Material online). In the one hand, SAR, response to SA, response to hypoxia, or response to chitin were activated in the evolved viruses, while amine metabolic process, cellular response to hypoxia and to cold, and cellular response to jasmonic acid stimulus were enriched in the group of DEGs repressed in the evolved viruses. In the other hand, some categories were found only in specific genotypes, *e.g*., cutin biosynthetic process, isoprenoid and tetrapyrrole metabolic processes were enriched within the DEGs activated by the viral adaptation in WT but not in the mutants. *ddm1* presented the largest number of specific terms within the DEGs activated in evolved lineages (*e.g*., carotenoid related processes, cellular homeostasis or ketone biosynthetic processes) while *jmj14* was the genotype with more specific terms within the DEGs repressed (*e.g*., cell wall biogenesis, sulfur compound biosynthetic process or tropism between the specifically enriched terms) (fig. 3B, supplementary fig. S2, Supplementary Material online).

### A core of genes associated with differences in TuMV adaptation was found differentially expressed in all plant genotypes

From a total of 9409 DEGs, we found a set of 1029 genes whose expression was significantly affected by the degree of TuMV adaptation, independently of the plant genotype in which evolution was carried on (fig. 3C). This core of genes showed a consistent profile of up- (48.0%) and down-regulation (51.5%) across all plant genotypes, except for five genes (fig. 3D). These exceptions were (fig. 3D, table S2, Supplementary Material online): AT1G69480 and AT4G09820 both specifically repressed in *dcl2 dcl3 dcl4* but activated in all other genotypes, AT2G40610 and AT5G65730 only repressed in *jmj14* but activated in the rest of genotypes, and AT4G25480 repressed in *dcl2 dcl3 dcl4*, *jmj14* and *polv* but activated in WT and *ddm1* plants.

The GO biological processes associated with the set of genes that increase their expression with virus adaptation in all genotypes include response to chitin, response to SA, SAR, response to ethylene signaling pathway and phosphorelay signal transduction system (fig. 3E and Supplementary fig. S2, Supplementary Material online). Among the shared genes whose expression has been repressed by viral adaptation, processes related with amine and aromatic amino acid family biosynthesis, jasmonic acid metabolic process, auxin metabolic processes, circadian rhythm, response to blue light, or response to toxic substances were observed (fig. 3E and supplementary fig. S2, Supplementary Material online). Categories such as cellular response to hypoxia, cellular response to ethylene stimulus, response to cold, or response to red or far-red light appeared enriched both in activated and repressed genes (fig. 3E and supplementary fig. S2, Supplementary Material online).

### Classification of genes according to their profile of gene expression in non-infected plants vs. plants infected with the ancestral or the evolved viruses

In order to get a wider picture of the effect of infection with viral lineages at different degrees of adaptation to their local host genotypes, we considered the DEGs across the three inoculation conditions (fig. 1 green lines) using the following approach. According to the inoculum type, *i.e.*, mock, ancestral and evolved viruses, genes can be classified into nine basic profiles based on the direction of the changes between each pair of conditions, as illustrated in fig. 4A. Fig. 4B shows the clustering of the five plant genotypes according to the similarity between the genes classified into each of the nine profiles. DNA methylation mutants *dcl2 dcl3 dcl4*, *ddm1* and *polv* cluster together (*polv* share 81% with *dcl2 dcl3 dcl4* and an 80% with *ddm1*) while the histone demethylation mutant *jmj14* shows a more dissimilar distribution of profiles.

**Figure 4.**
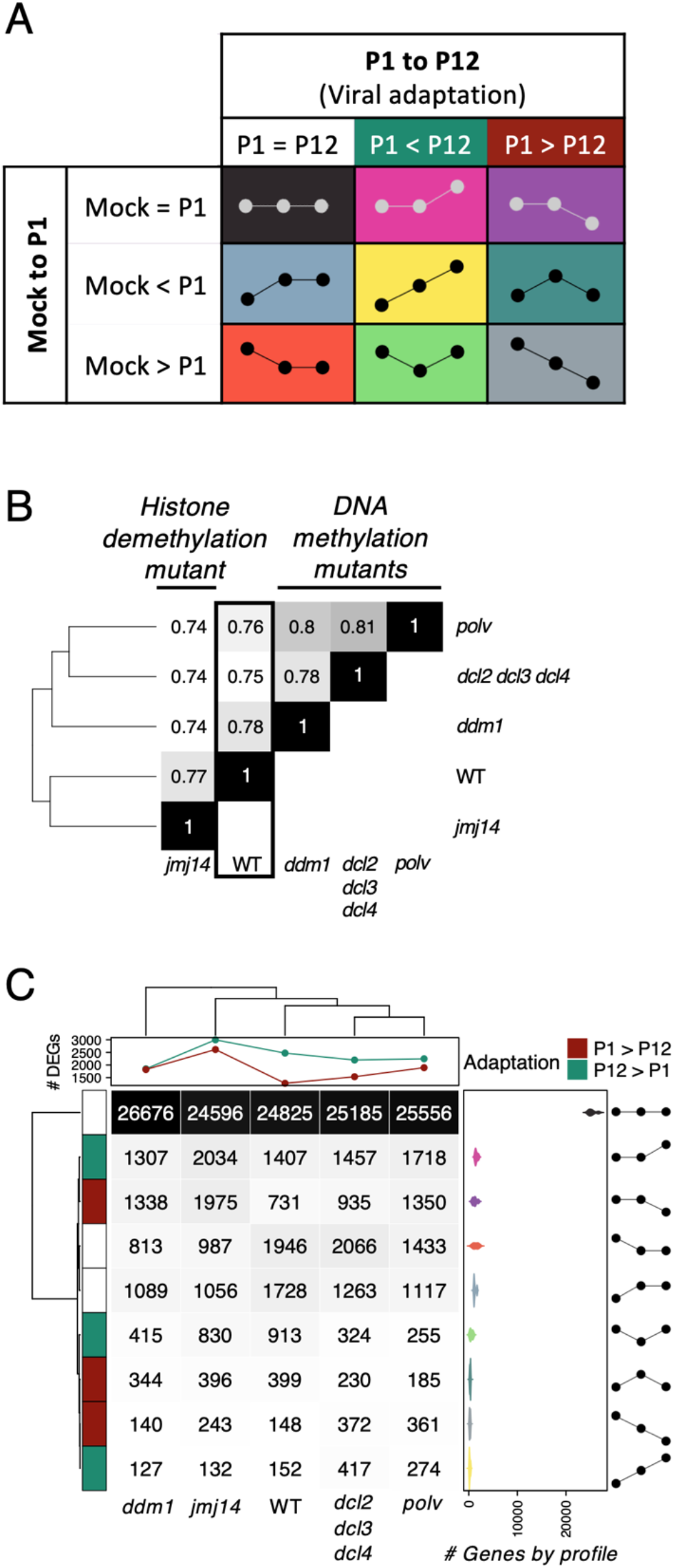
Classification of gene expression profiles according to the nine possible patterns of variation across conditions. (A) Classification profiles based on the direction of changes between the three inoculation conditions, mock-inoculated, ancestral (P1) virus-infected and evolved (P12) viruses-infected. (B) Hierarchical clustering of plant genotypes based on the similarity amongst their classification profiles in panel (A). (C) Abundance of each profile per plant genotype.

Fig. 4C shows all DEGs found on each of the plant genotypes classified into each of the nine possible expression patterns when comparing between mock-inoculated, ancestral- and evolved-infected plants. By far, in all five genotypes the most common pattern corresponded to genes that showed no differences between mock-inoculated and virus-infected plants, followed in WT by genes that responded to infection in the same magnitude (up or down) both for the ancestral and the evolved viruses. The most interesting cases are precisely those in which the pattern showed differences between WT and mutant plants inoculated with ancestral and evolved viruses. Fisher’s exact tests were used to evaluate pairwise differences between the patterns of DEGs shown for WT and each of the four mutant plant genotypes. In all four cases, the number of DEGs up or down found in the ancestral and the evolved inoculated mutant plants was different than observed for WT plants (Fisher’s exact test, *P* < 0.001 in all comparisons). Indeed, the number of up- and down-regulated DEGs due to virus evolution relative to the number of DEGs not associated to virus evolution (fig. 4C) is 2.75-fold largest for *jmj14* followed by 1.93-fold for *ddm1*, while the smallest difference was found for WT plants (1.02-fold). This observation further supports that TuMV adaptation to histone demethylation mutant plants resulted in a stronger genome-wide alteration in gene expression than adaptation to WT plants.

Next, we sought to functionally categorize the DEGs specific to each of the nine patterns in fig. 4C. Supplementary fig. S3, Supplementary Materials online, present the results of the functional enrichment GO analyses per pattern for each plant genotype. For example, comparing across plant genotypes, a pervasively enriched GO term was photosynthesis processes though classified into different profiles: downregulated upon infection in *dcl2 dcl3 dcl4*, *polv* and WT, downregulated only by evolved infection in *jmj14* and upregulated in evolved-infected *ddm1* plants. Responses to chitin also showed an upregulation pattern for *ddm1* plants infected with the evolved viral lineages but not in the other genotypes. As an additional example of highly variable profiles, responses to SA signaling were always upregulated upon infection with evolved viruses, except in *ddm1* plants, and with *polv* plants showing upregulation also when infected with the ancestral viruses.

To delve deeper into functional responses potentially influenced by virus adaptation, we specifically examined each plant genotype and focused on differences between virus ancestral- and evolved-infected plants (supplementary fig. S4, Supplementary Material online). In WT plants, only responses to cold were significantly altered by both types of viral lineages. Processes related to O_2_ levels and SA were overrepresented in evolved-infected plants, while processes related to membrane transport and organization, endocytosis, rRNA processing, and cytoplasmic translation were underrepresented in evolved-infected plants. In *dcl2 dcl3 dcl4*, responses to chitin, and hypoxia were specifically overrepresented in evolved-infected plants, while processes related to circadian rhythms and the transition from vegetative to reproductive phases in meristems were underrepresented in the same plants. This observation aligns with the impact of TuMV infection on the transition from vegetative growth to reproduction, resulting in plant castration (Melero et al., 2023). *ddm1* plants exhibited the shortest list of significant GO terms, with only photosynthesis being shared by three of the expression profiles, two of which were overrepresented in evolved-infected plants compared to their ancestral counterparts. The histone modification-deficient *jmj14* plants displayed a diverse array of GO terms across the nine expression patterns. For example, responses to chitin, hypoxia, SA, jasmonic acid, and SAR were overrepresented in evolved-infected plants compared to ancestral-infected ones. In contrast, photosynthesis, response to cold, biogenesis of ribonucleoprotein complexes, rRNA processing and metabolism, and, notably, brassinosteroid-mediated signaling were underrepresented among evolved-infected plants. Finally, in *polv* plants, plastids organization and microtubule-related processes were specifically overrepresented in evolved-infected plants, while endomembrane system organization, Golgi vesicle transport, and processes related to the transition from vegetative to reproductive phases in meristems and regulation of multicellular differentiation were significantly underrepresented among evolved-infected plants compared to ancestral-infected ones. In short, an analysis of the intersections by profile shows that, avoiding the no-changes profile, the pattern with more genes in common among genotypes was represented by the repression in plants infected with ancestral viruses but not change with evolution (242 genes in common) while the less convergent was the upregulation through all conditions (6 genes in common) (supplementary fig. S5, Supplementary Material online).

### DNA but not histone methylation affects the expression of genes nearby TEs

In a prior study, Corrêa et al. (2020) observed that in WT plants infected with evolved TuMV lineages, but not with their ancestral counterparts, TEs concentrated in centromeric and pericentromeric regions were induced early after infection. At later stages of infection, both induction and repression of different TE families were observed. Subsequently, Corrêa et al. (2024) demonstrated that POLV target genes had a higher percentage of TE marks around their transcription start sites (TSS) than other genes. In contrast, genes regulated by JMJ14 showed a lower level of marks around their TSS. These observations suggest that genes found to be upregulated in infected DNA methylation mutants, especially in *ddm1*, but less in *jmj14* plants, are likely to be controlled by TE-related mechanisms. To further investigate this hypothesis, we sought differences in the number of DEGs with a TE within 1 Kbp upstream of their TSS between virus ancestral- and evolved-infected plants (fig. 5A). First, a highly significant correlation exists between the total number of DEGs and the number of DEGs in the proximity of TEs (fig. 5A: Pearson’s *r* = 0.990, 3 d.f., *P* = 0.001). The proportion of DEGs associated with TE was slightly higher in all the mutants than in WT plants, but this difference was significantly enriched only in *ddm1* plants (fig. 5A: Fisher’s exact test, *P* = 0.045). However, when comparing the TE-related DEGs between the three DNA methylation mutants and the histone modification mutant *jmj14*, also *ddm1* showed a significant augment in the number of TE-associated DEGs responding differentially to ancestral and evolved viral populations (fig. 5A: Fisher exact test, *P* = 0.036). The observed differences in the number of DEGs close to TEs in *ddm1* plants could be attributed to the higher basal instability of these genetic elements and the stronger transcriptome disturbance caused by the evolved virus (Supplementary fig. S1B, Supplementary Material online) in this genotype compared to the others.

**Figure 5.**
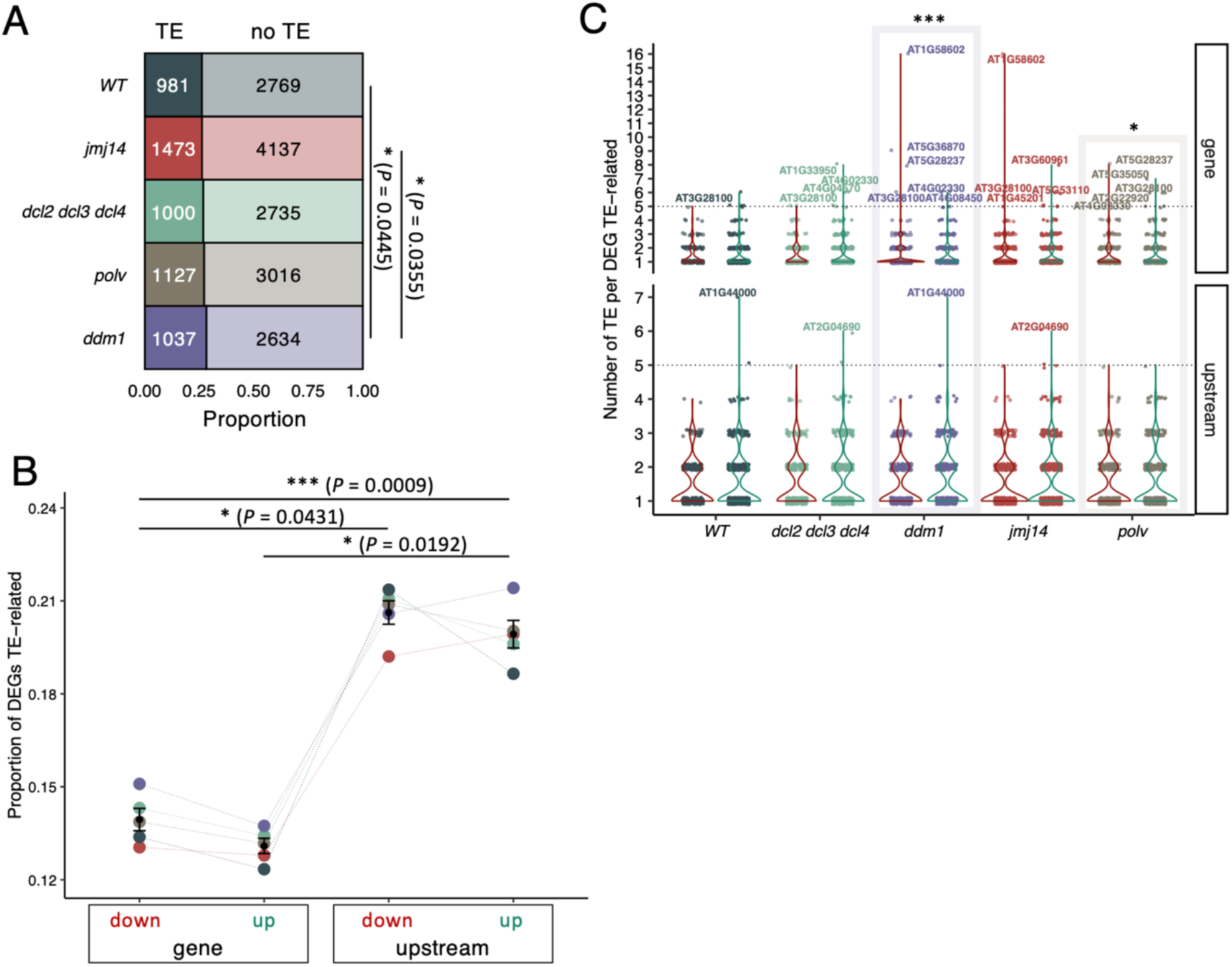
Analysis of DEGs (contrasting evolved *vs*. ancestral viruses) in the context of their association to TEs. (A) Proportion of DEGs in close proximity to TEs in each plant genotype. *ddm1* present a higher number of DEGs TE-associated than WT and *jmj14*. (B) Interaction between the position of TEs in genes (inside or within 1 Kbp upstream of their TSS) and the direction of regulation (upregulated or downregulated in the evolved lineages). Values are normalized by the total number of DEGs with the same locus and regulation. *P* values of Kruskall-Wallis rank sum test are indicated over the significant contrasts. (C) Distribution of DEGs related to the presence of TEs. Violin plots color represents DEGs-TE related within the activated (green) and repressed (red) ones. *ddm1* and *polv* showed significant differences in the distribution of its TEs depending on the localization and the expression where the TEs are associated (Kruskal-Wallis, *P* < 0.001 and *P* = 0.001, respectively).

Subsequently, we categorized the DEGs close to TEs based on their regulation (upregulated or downregulated in the evolved lineages) and the position of the TEs in the gene (whether they were located inside or within 1 Kbp upstream TSS) (fig. 5B). Significant differences exist between the number of DEGs containing a TE inside or upstream the gene (Kruskal-Wallis test: χ^2^ = 6.818, 1 d.f., *P* < 0.001), the sign of the regulation of the TE-related DEGs (Kruskal-Wallis test: χ^2^ = 4.811, 1 d.f., *P* = 0.028) and the interaction between both TE location and the sign of regulation (Kruskal-Wallis test: χ^2^ = 16.006, 3 d.f., *P* = 0.001). A Dunn *post hoc* pairwise comparisons test found significant differences between the location of TEs (in down-regulated, *P* = 0.043; in up-regulated, *P* = 0.019) but the regulation component only was significant between different locus (*P* < 0.001) (fig. 5B). No significant differences between genotypes were found in any group. The data suggest that TEs near gene promoters are more likely to correlate with changes in gene expression than those within genes.

Additionally, we checked the number of TEs associated with each TE-related DEG. This data revealed that although the proportion of DEGs having TEs in proximity of TSS is higher than the ones having TEs inside the genes (fig. 5B), the total counts of TEs in these same DEGs is higher inside than upstream the genes (fig. 5C: Wilcoxon rank test with continuity correction, *P* < 0.001). After separating by sign of regulation and localization, *ddm1* (Kruskal-Wallis, *P* < 0.001) and *polv* (Kruskal-Wallis, *P* = 0.001) presented differences within its four possible combinations (fig. 5C). The DEG with the highest density of TEs found inside its sequence (16 of both classes of TE) was *RECOGNITION OF PERONOSPORA PARASITICA 7* (*RPP7*), a gene that encodes an LRR and NB-ARC domains-containing disease resistance protein (fig. 5C) (Liu et al. 2015). *RPP7* appeared as repressed by the evolved viruses adapted in *ddm1* and *jmj14* mutant plants. This gene is known to be regulated by a particularly interesting epigenetics mechanism in which a “domesticated” TE located in its first intron must be properly silenced in order to produce full-length transcripts (Tsuchiya and Eulgem 2013). Out of these 16 TEs inside *RPP7*, half belonged to the MuDR superfamily (fig. 6) and more precisely to the VANDAL transposons. VANDAL transposons have coevolved mechanisms to escape the epigenetic silencing (Sasaki et al. 2022). It is noteworthy that in *jmj14* and WT plants both activated and repressed genes were enriched in MuDR inside coding regions (fig. 6). However, in *polv* plants only activated genes were enriched in MuDR while in *ddm1* plants, MuDR enrichment was only for repressed genes.

**Figure 6.**
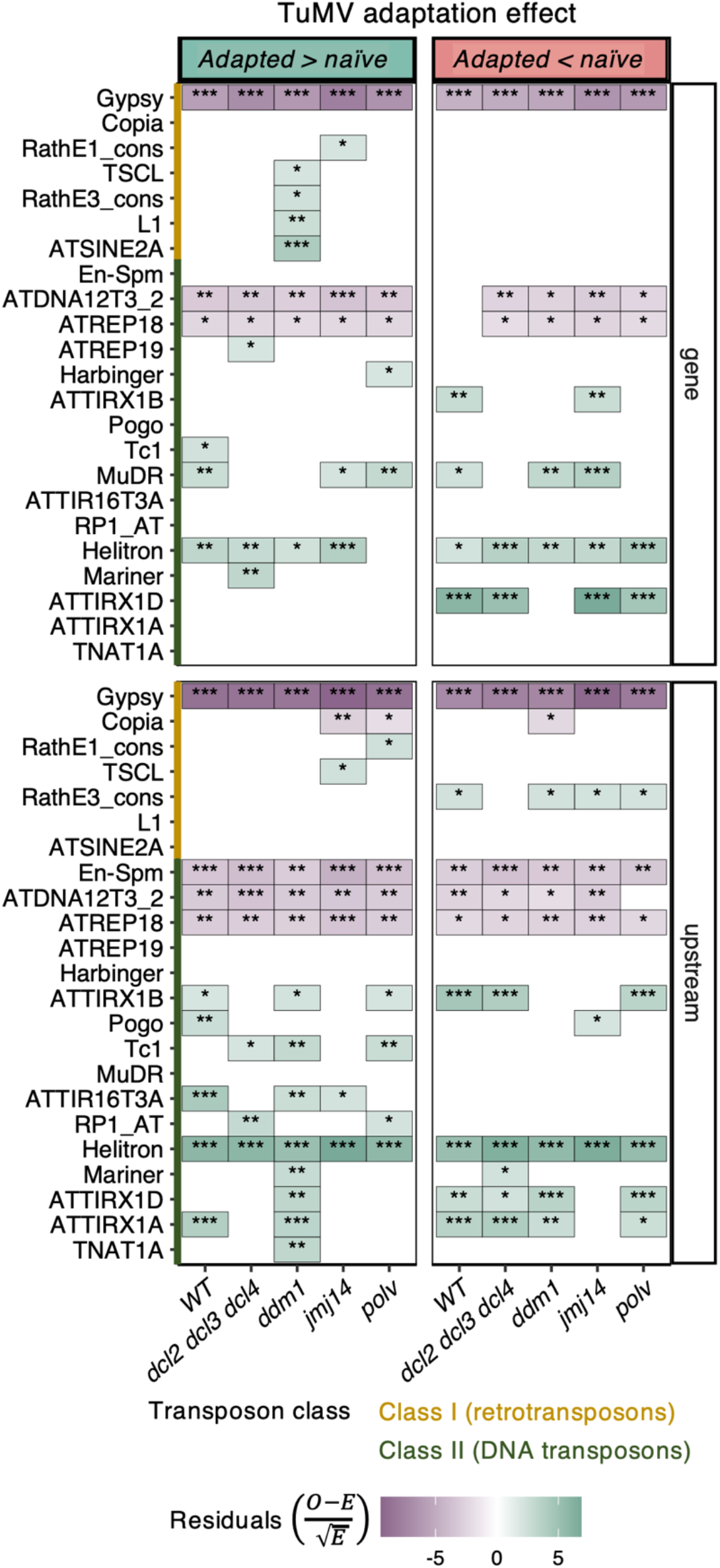
Analysis of the TE families linked DEGs associated to differences between ancestral and evolved viral isolates. Asterisks indicate significance levels in a χ^2^ goodness-of-fit test: * *P* < 0.05, ** *P* < 0.001 and *\*\*\* P* < 0.001.

### Class II TEs are significantly enriched among Arabidopsis genes responding to TuMV adaptation while class I TEs were significantly deprived

An enrichment analysis of the types of TEs associated with the DEGs associated with virus adaptation revealed a general pattern of overrepresentation of class II elements (DNA transposons) and underrepresentation of class I ones (retrotransposons). A detailed analysis by superfamilies of TEs is shown in fig. 6. The distribution of TE families across all the genome was used as the null model (expected values) and the proportion found at each condition (observed values) was tested with a χ^2^ goodness-of-fit. Briefly, from class I, Gypsy appears less than expected (*P* < 0.001), no matter the sign of regulation, localization relative to the gene coding sequence, or the plant genotype. An interesting exception is *ddm1* plants, where ATSNE2 (SINE type) and L1 (LINE type) were significantly enriched within the activated genes (fig. 6: *P* < 0.001 and *P* = 0.008, respectively).

In contrast with this deprivation in class I TEs, a general enrichment in class II transposons was observed in all four plant genotypes, both upstream the TSS and inside the DEGs (*P* < 0.001). Members of the Helitron superfamily were the most abundantly over-represented type (fig. 6). Helitrons (alike ssDNA viruses, bacteria and plasmids) use rolling circle replication for its propagation. This mechanism, in contrast with the cut-and-paste strategy of other DNA transposons, allows the retention of the original TE while the daughter TEs is copied. This mechanism allows TEs to capture gene fragments that can derive in epigenetic conflict (Coates 2015; Barro-Trastoy and Köhler 2024). Cases of epigenetic conflict in the stress response context driven by Helitrons have been previously reported. For example, ATREP2 TEs have been found enriched in genes regulating ISR responses to herbivores (Wilkinson et al. 2023; Barro-Trastoy and Köhler 2024).

A GO enrichment analysis of the biological processes of these TE-proximal DEGs (supplementary fig. S7A, Supplementary Material online) showed that the response to SA appeared overrepresented in all five plant genotypes. The same occurs with various categories related to hypoxia and the response to chitin, represented in genes related to both classes of TE. Responses to ethylene and to cold were also overrepresented in at least four of the five plant genotypes (supplementary fig. S7A, Supplementary Material online). *ddm1* plants presented the largest number of terms enriched within the DEG TE-related elicited by the viral adaptation and none within the repressed ones (supplementary fig. S7B, Supplementary Material online). Its more specific categories are related to photosynthesis. In contrast, *polv* present the largest number of enriched terms within the TE-related DEGs repressed in the evolved viruses (supplementary fig. S7B, Supplementary Material online). In this case, genes are related to class I TE and all the functional categories are related to metabolic processes (supplementary fig. S7A, Supplementary Material online).

### Putative alteration of the TuMV-*A. thaliana* protein-protein interaction network due to virus adaptation

The way in which virus and host interact changes with the degree of virus’ adaptation, which indeed depends on the fine tuning in the interactions between viral and host factors (Agudelo-Romero et al. 2008; Cervera et al. 2018). In order to explore the extent to which the target genes of TuMV adaptation could be perturbing the virus-host interaction network, we integrated the transcriptomic results described above with the protein-protein interaction (PPI) data published by Martínez et al. (2023).

Fig 4A depicts expression profiles for DEGs whose proteins have been demonstrated to interact with viral proteins. The nine derived profiles differ significantly from the genome-wide expression in four host genotypes, except for the *polv* mutant [χ^2^ homogeneity tests; *dcl2 dcl3 dcl4* (*P* < 0.001), *ddm1* (*P* = 0.011), *jmj14* (*P* = 0.002), *polv* (*P* = 0.271), and WT (*P* = 0.045)] (fig. 7A). Focusing on the DEGs showing differences between virus ancestral- and evolved-infected plants, we found in infected *ddm1* and WT plants more PPI-associated DEGs activated within the viral adaptation, while *dcl2 dcl3 dcl4* and *jmj14* were enriched in more repressed genes (fig. 7B). We used the number of genes activated and repressed by the evolved viral lineages as a proxy of the probabilities for gene activation and repression in each host genotype. With these probabilities, we calculated the expected number of interactors altered for each viral protein (fig. 7C) and used a χ^2^ homogeneity test to evaluate whether some of the viral proteins were enriched or depleted in selected DEGs. P3 and P3N-PIPO showed more interactions than expected with proteins encoded by the overexpressed genes in *ddm1* (χ^2^ homogeneity tests, *P* < 0.001 for both proteins), an over-representation of repressed genes was found for 6K1 in WT (*P* = 0.031), for CI and NIb in *dcl2 dcl3 dcl4* (*P* = 0.029 and *P* = 0.007, respectively), as well as for CI in *jmj14* (*P* = 0.021). Despite this, *ddm1* is the only genotype with significantly more upregulated interactors than expected by share chance.

**Figure 7.**
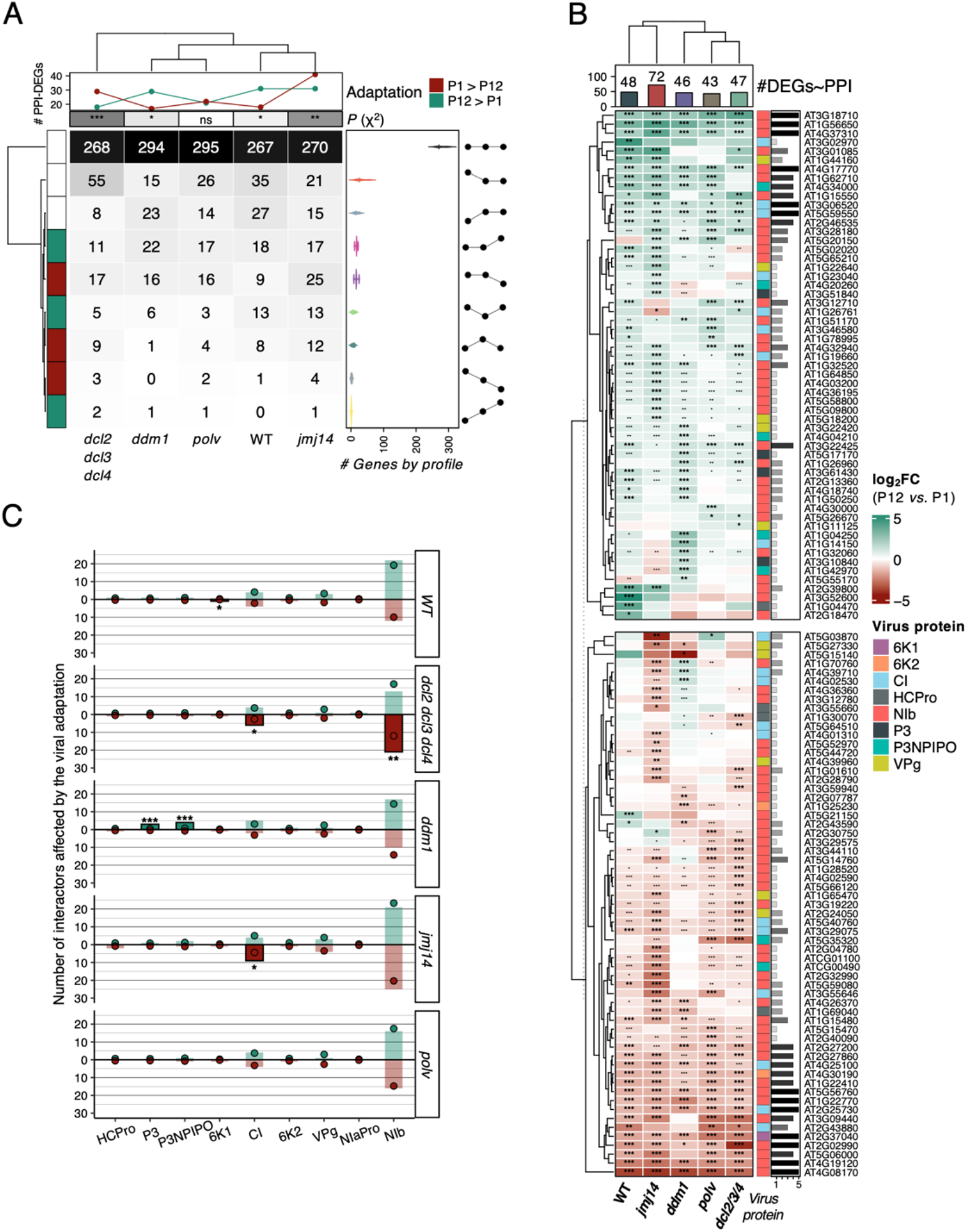
Interactors in the host-virus PPI network are altered by the adaptation of the virus. (A) Distribution of the PPI-related genes in the nine profiles classification developed in this work. The expression profiles (fig. 4A) of the genes involved in the PPI, were found to significantly differ from the genome-wide expression in four host genotypes, except in the case of *polv* mutant [χ^2^ homogeneity tests; *dcl2 dcl3 dcl4* (*P* < 0.001), *ddm1* (*P* = 0.011), *jmj14* (*P* = 0.002), *polv* (*P* = 0.271) and WT (*P* = 0.045)]. (B) Gene expression of DEGs PPI-associated. Asterisks indicate significant changes (adjusted *P* < 0.05), when |log_2_*FC*| > 1, bolded. The lateral bars represent the number of genotypes with the interactor significantly affected (adjusted *P* < 0.05 and |log_2_*FC*| > 1). (C) Observed (bars) and expected (points) host interactors for each viral protein were tested to search for significant differences using a χ^2^ homogeneity test. Asterisks indicate significance level: * *P* < 0.05, ** *P* < 0.001 and *\*\*\* P* < 0.001. P1: ancestral viruses, P12: evolved viruses.

Finally, supplementary fig. S8, Supplementary Material online, maps the effects of viral adaptation in the host transcriptome over the host-virus PPI network. Thirteen interactors were found altered in all five plant genotypes (supplementary table S3, Supplementary Material online). Viral replicase NIb had five repressed (with diverse functions, but remarkably a methyltransferase superfamily member) and four activated interactors (including transcription factor MYB75 and and E3 ubiquitin ligase), multifunctional CI two activated (E3 ubiquitin ligase and an Agenet domain-containing protein involved in chromatin remodeling) and one repressed (Zinc finger FYVE domain containing protein), and viroporin 6K1 a repressed one, *PHENYLALANINE AMMONIA-LYASE 1* (*PAL1*). Although these results merit further exploration, we refer the readers to the information displayed in table S3, Supplementary Material online, while focusing here only in *PAL1*.

Pentamers of 6K1 had been demonstrated to be a functional viroporin (Chai et al. 2024). Viroporins are small hydrophobic proteins that insert in membranes modifying its permeability. Their role in viral pathogenicity and inflammatory responses has been assessed in several viruses (Nieva, Madan and Carrasco 2012), as well as their potential as drug targets (To, Surya and Torres 2016). 6K1, contrary to the majority of the known viroporins, also plays an essential role in the viral replication cycle (Cui and Wang 2016). 6K1 interactor PAL1 is a phenylalanine ammonia-lyase that acts upstream of the phenylpropanoid pathway. This pathway was found to be consistently repressed in plants infected with the evolved TuMV populations (fig. 3 and Supplementary fig. S3, Supplementary Material online), thus suggesting its importance in viral adaptation. Among other diverse roles, this pathway has been shown to be redundant in synthetizing SA and to be involved in resistance to fungal (Ferrari et al. 2003) and viral (Kavil et al. 2021) pathogens. Concretely, the overexpression of *PAL1* in cassava increases resistance to cassava brown streak virus, another member of the *Potyviridae* family (Kavil et al. 2021). Despite the character of the interaction 6K1 - PAL1 remains unknown, the repression of *PAL1* expression by the evolved lineages in the five plant genotypes suggests *PAL1* as a promising candidate for future studies.

### Concluding remarks

Evolution deeply changes the way plant viruses interact with their hosts (Agudelo-Romero et al. 2008; Hillung et al. 2016; Corrêa et al. 2020). Following up with previous results (Ambrós et al. 2024), the comparative analyses of large transcriptomic datasets have shown that alterations in epigenetic regulatory pathways modulate the evolution of RNA viruses. We have identified a common set of genes that affect TuMV adaptation to *A. thaliana* regardless of the specific genetic makeup of the plant’s epigenetic pathways, but also many genes whose effect on viral adaptation was host-genotype dependent. Our results reinforce the idea that *jmj14* histone demethylase represents a major driving factor during TuMV evolution. Our analyses also discovered a significant association between genes regulated by epigenetic marks and the presence of TEs within or upstream the genes involved in TuMV adaptation, especially in the *ddm1* plants. Finally, integrating transcriptomic data with PPIs in the same pathosystem has allowed us to identify additional host genes involved in TuMV adaptation, highlighting the role of the interaction between the viroporin 6K1 and PAL1 in virus adaptation.

## Materials and Methods

### Plants, virus and experimental evolution

Full details of the evolution experiment can be found in Ambrós et al. (2024). In short, in addition to the wild-type *A. thaliana* (L.) HEYNH Col-0 accession, four mutants have been used in this study: *dcl2 dcl3 dcl4*, *ddm1* and *polv* affecting RdDM, and the *jmj14* histone demethylase. In all experiments, plants were maintained in a climatic chamber under a photoperiod of 8 h light (LED tubes at PAR 90 - 100 μmol m^−2^ s^−1^) at 24 °C and 16 h dark at 20 °C and 40% relative humidity, in a mixture of 50% DSM WNR1 R73454 substrate (Kekkilä Professional, Vantaa, Finland), 25% grade 3 vermiculite and 25% 3 - 6 mm perlite. Pest management was performed by the introduction of *Stratiolaelaps scimitus* and *Steinernema feltiae* (Koppert Co., Málaga, Spain).

TuMV infectious sap was obtained from TuMV-infected *Nicotiana benthamiana* DOMIN plants inoculated with the infectious plasmid p35STunos containing a cDNA of the TuMV genome (GenBank AF30055.2) as described in Corrêa et al. (2020). This sequence variant corresponds to the YC5 strain from calla lily (Chen et al. 2003). Symptomatic tissues were pooled, frozen with liquid N_2_ and homogenized using a Mixer Mill MM400 (Retsch GmbH, Haan, Germany).

Five TuMV lineages were evolved during twelve consecutive serial passages in each one of the six selected mutant genotypes (fig. 1). To begin the evolution experiment, ten *A. thaliana* plants per lineage and genotype were gently rubbed onto two leaves with 5 μL of inocula [0.1 g of the above stock diluted in 1 mL inoculation buffer (50 mM phosphate buffer pH 7.0, 3% PEG6000, 10% Carborundum)]. Plants were all inoculated when growth reached stage 3.5 in the Boyes et al. (2001) scale. This synchronization ensures that all hosts were at the same phenological stage when inoculated. The next passages were made by harvesting the symptomatic plants at 14 dpi, preparing the infectious sap as described above, and inoculating it into a new healthy batch of ten plants.

### Total RNA extractions and preparation of samples for HTS

Pools were made of 10 infected symptomatic plants per lineage, genotype, and serial passage, frozen with liquid N_2_ and preserved at −80 °C until it was homogenized into fine powder. Aliquots of ∼0.1 g were used for total RNA (RNAt) extractions with the Agilent Plant RNA isolation Mini kit (Agilent Technologies, Santa Clara CA, USA). Three aliquots of total RNA (RNAt) per sample were separated and their concentrations adjusted to 50 ng μL^−1^.

For RNA-Seq, RNAt from 70 - 90 mg infected and healthy plants was extracted using the GeneJET Plant RNA Purification Mini Kit (Thermo Fisher Scientific, Waltham MA, USA) following the manufacturer’s instructions. RNA quality was checked with NanoDrop One (Thermo Fisher Scientific) and agarose gel electrophoresis and integrity and purity tested with Bioanalyzer 2100 (Agilent Technologies). Libraries preparation and Illumina sequencing was done by Novogene Europe using a NovaSeq 6000 platform and a Lnc-stranded mRNA-Seq library method, ribosomal RNA depletion and directional library preparation, 150 paired end, and 6 Gb raw data per sample. Novogene did quality check of the libraries using a Qubit 4 Fluorometer (Thermo Fisher Scientific), qPCR for quantification and Bioanalyzer for size distribution detection.

### HTS data processing

The quality of the Fastq files was assessed with FASTQC (http://www.bioinformatics.babraham.ac.uk/projects/fastqc/) and MultiQC (Ewels et al. 2016) and preprocessed as paired reads with BBDuk (https://sourceforge.net/projects/bbmap/). Adapters were removed, the first ten 5’ nucleotides of each read were cut, and the 3’ end sequences with average quality below 10 were removed. Processed reads shorter than 80 nucleotides were discarded. The parameter values were set to: *ktrim* = r, *k* = 31, *mink* = 11, *qtrim* = r, *trimq* = 10, *maq* = 5, and *forcetrimleft* = 10, and *minlength* = 80. The processed files were mapped to TuMV isolate YC5 with the BWA-MEM algorithm (Li 2013). Resulting SAM files were binarized and sorted with SAMtools (Danecek et al. 2021) and duplicates marked with the MarkDuplicates method of GATK version 4.2.2.0 (McKenna et al. 2010).

Processed files were mapped to the *A. thaliana* genome (TAIR10) with STAR version 2.7.9a (Dobin et al. 2013), using the run mode ‘alignReads’ and the quantification mode ‘GeneCounts’. The quantification results were used for the transcriptome analysis. Resulting SAM files were then binarized and sorted with SAMtools (Danecek et al. 2021). All the R and shell codes generated for this project are available in https://github.com/MJmaolu/evolution_TuMV_in_A.thaliana_Epigenetic_Mutants/tree/main.

### Host gene expression and functional annotation

The counts matrix was constructed from the STAR results (reversed-stranded column). The three blocks of differential expression analyses (Supplementary fig. S4, Supplementary Material online) were performed using DESeq2 v1.36.0 (Love, Huber and Anders 2014), considering as significant genes those with an adjusted *P* < 0.05 (Benjamini and Hochberg method) and a |log_2_*FC*| > 1. A first block of contrasts (mutants *vs*. WT samples by condition, bourgogne lines in fig. 1) was carried out to study the differences in the responses of each mutant genotype relative to the responses of the WT plants. This contrast was applied to the mock-inoculated samples, ancestral virus-inoculated samples and, evolved virus-inoculated samples separately. A second block of contrast [adapted *vs.* naïve infection by genotype, purple differential expression analyses (DEAs) in fig. 1] the focus was on identifying DEGs between samples infected with an evolved virus *vs*. those infected with the ancestral virus, for which we considered samples infected by the same viral lineage as paired samples. For the third group of analysis (expression across three inoculation conditions by genotype, green lines in fig. 1) an additional group of DEAs to identify the differences between naïve infection and mock by each genotype was performed. The classification method is described below. For functional enrichment analyses, STRINGdb v2.8.4 (Szklarczyk et al., 2021) and clusterProfiler v4.4.4 (Wu et al., 2021) were used.

All resulting DEGs were screened for the presence of annotated TEs within their ORFs or up to one thousand nucleotides upstream of their promoters using the GenomicRanges package, v1.48.0 (Lawrence et al. 2013) and the *A. thaliana* transposon annotation file from TEtranscripts (https://labshare.cshl.edu/shares/mhammelllab/www-data/TEtranscripts/TE_GTF/, last accessed 01/26/2023). Functional enrichment analysis of the resulting genes was performed as previously described.

All the processes were conducted with home-made scripts in R (version 4.2.1), available in the repository mentioned above where a reproduction of the session information can also be found.

### Classification of gene expression across mock-ancestral viruses-evolved viruses into nine basic profiles

All the genes were classified in the nine possible profiles (fig. 3A) depending on the change in the expression between the three inoculation conditions. For evaluating the direction of changes (mock *vs.* ancestral and ancestral *vs.* evolved) the results obtained with DESeq2 for each genotype were used. For each of both changes, cases were assigned into three categories: (*i*) increasing expression if adjusted *P* < 0.05 and log_2_*FC* > 1, (*ii*) decreasing expression if adjusted *P* < 0.05 and log_2_*FC* < –1 and (*iii*) non-affected otherwise.

### PPI network and viral protein enrichment

The *A. thaliana* – TuMV PPI published by Martínez et al. (2023) was used. The DEGs selected in the contrast evolved *vs.* ancestral (purple lines in fig. 1) were represented in the PPI. Node sizes indicate the number of genotypes where the gene is significantly altered (adjusted *P* < 0.05) with a |log_2_*FC*| > 1. The network was generated with Cytoscape version 3.10.0 (Shannon et al. 2003). We used the number of genes activated and repressed by the adapted viral lineages as a proxy of the probabilities for gene activation and repression in each host genotype. With these probabilities, we calculated the expected number of interactors altered for each viral protein (fig. 7C) and used a χ^2^ homogeneity test to evaluate whether some of the viral proteins were enriched or depleted in selected DEGs. Significant changes were considered when *P* < 0.05.

## Supplementary Material

Supplementary data are available at *Molecular Biology and Evolution* online.

**Supplementary fig. S1**. (A) Common DEGs to all mutant plants mock-inoculated, infected with ancestral (P1) or with evolved (P12) viruses compared to WT plants with an opposite regulation. Color and numbers indicate log_2_*FC* values (mutant *vs.* WT). (B) Change in the number of total and genotype-specific DEGs between mutant plants mock-inoculated, infected with the ancestral P1 or with the evolved P12 compared to WT plants. Lines indicate the percentage of specific DEGs over the total.

**Supplementary fig. S2**. Relates to fig. 3A. Functional enrichment (GO: Biological Processes) analysis of DEGs for the five plant genotypes. Categories common across all plant genotypes, shared by more than one genotype or specific of a particular genotype are indicated by different symbols. Symbol colors represent whether the DEGs are enriched (green) or under-represented (red) in plants infected with evolved (P12) *vs.* ancestral (P1) viruses. Only categories with adjusted *P* < 0.001 are shown.

**Supplementary fig. S3**. Relates to fig. 4. Functional enrichment (GO: Biological Processes) analysis of DEGs classified according to the nine expression profiles.

**Supplementary fig. S4**. Shared DEGs per expression profile and plant genotype. **Supplementary fig. S5**. Functional enrichment (GO: Biological Processes) analysis of DEGs that are specific of each plant genotype. Each expression profile is represented in a different panel.

**Supplementary fig. S6**. Functional enrichment (GO: Biological Processes) analysis of DEGs. (A) Shared by all mutants and according to the expression profile. (B) Shared only the by the three mutants involved in DNA methylation pathway.

**Supplementary fig. S7**. (A) Functional annotation (GO: Biological Processes) of DEGs according to their relation with TEs. Open symbols represent class I TEs-DEGs associated and solid symbols represent class II TEs-DEGs associated. Green symbols represent DEGs activated while red symbols represent DEGs repressed in plants infected with the evolved (P12) viruses compared to plants inoculated with their (P1) counterparts. (B) Number of GO terms per plant genotype by regulation and TE-DEG type associated as in (A).

**Supplementary fig. S8**. *A. thaliana* – TuMV PPI network affected by the viral adaptation. Viral proteins are indicated by hexagons of different colors as in fig. 7B. Host proteins are indicated by circles. The size of the circle is proportional to number of genotypes where the gene is altered from none (smaller) to all (bigger node). The change in gene expression between plants infected with the ancestral (P1) or the evolved (P12) viruses is indicated by the node color, red if repressed by viral adaptation (P1 > P12) and green if activated (P12 > P1). Darker border indicates the gene expression is already altered in WT.

**Supplementary table S1**. Genes with a differential gene expression in all the mutants respect WT under the same inoculation conditions.

**Supplementary table S2**. Genes with opposed regulation within the core of DEGs involved in TuMV adaptation to *A. thaliana*.

**Supplementary table S3.** Genes of the PPI network affected by viral adaptation to all plant genotypes. When circles contain both colors, it means that the sign of the expression varied among host genotypes.

## Data availability

Raw Illumina RNA-Seq data generated for this study are available in NCBI SRA under BioProject accession PRJNA974369.

## Author contributions

S.F.E. conceived the experiments. S.A. and R.L.C. performed the experiments. M.J.O-U., R.L.C., S.F.E. conceived the bioinformatic analysis. M.J.O-U. wrote the code, performed all bioinformatic analyses and create the visualizations. M.J.O-U. and S.F.E. performed the statistical analyses. S.F.E. and M.J.O-U. wrote the first version of the manuscript. All authors edited and approved the latest version of the manuscript.

## Acknowledgements

We thank Francisca de la Iglesia and Paula Agudo for excellent technical support and the EvolSysVir lab members for comments and fruitful discussions. Computations were performed on the Garnatxa HPC cluster at the Institute for Integrative Systems Biology (I2SysBio).

