## Supplementary Material online for "Transcriptomic Insights into the Epigenetic Modulation of Turnip Mosaic Virus Evolution in *Arabidopsis thaliana*"

A Common DEGs (mutant *vs* WT) with opposite regulation across mutants

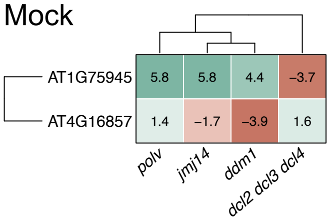


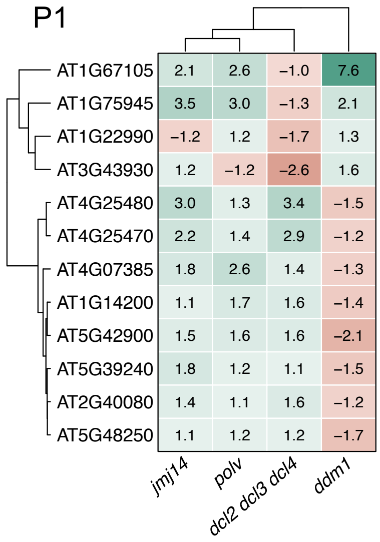


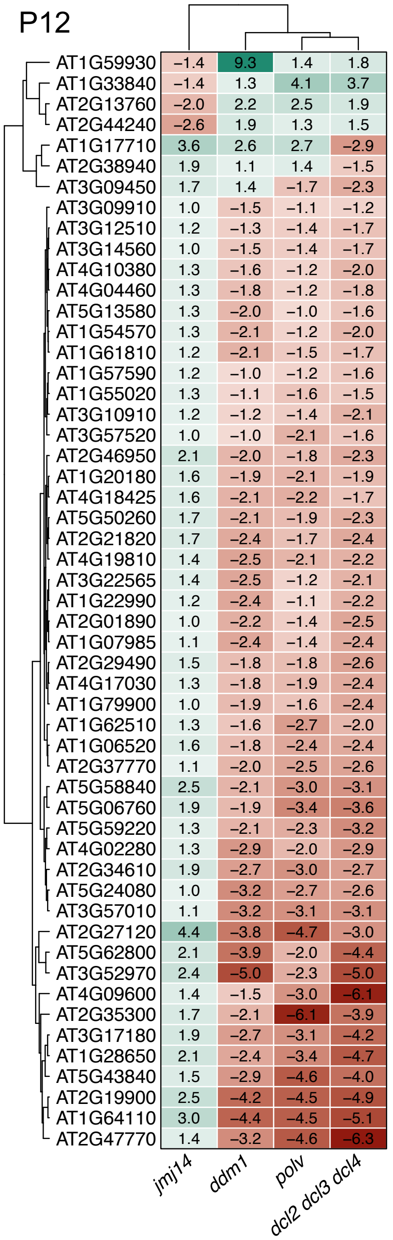


B Number of total and specific DEGs by mutant (*vs.* WT)

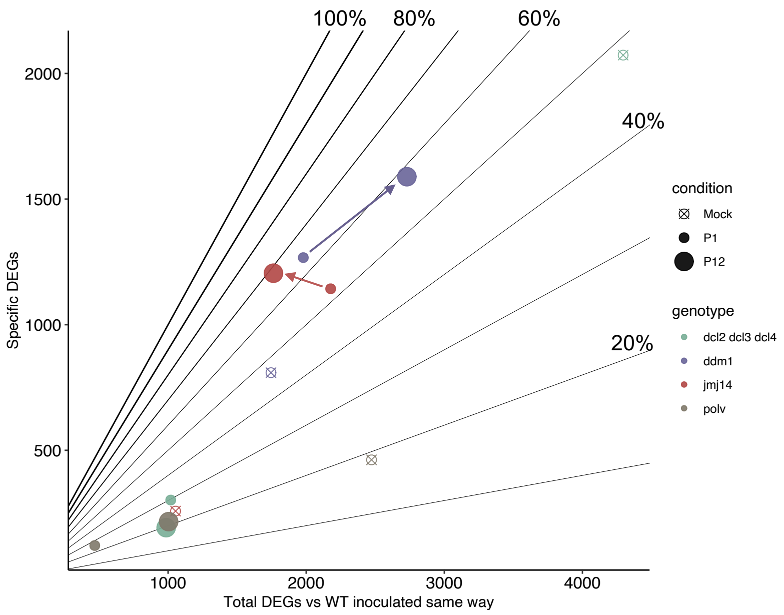


Supplementary Fig. S1

Functional enrichment analyses. GO: Biological processes separated by the direction of the change and its genotype specificity (Bonferroni-Hochberg adjusted $P < 0.001$).

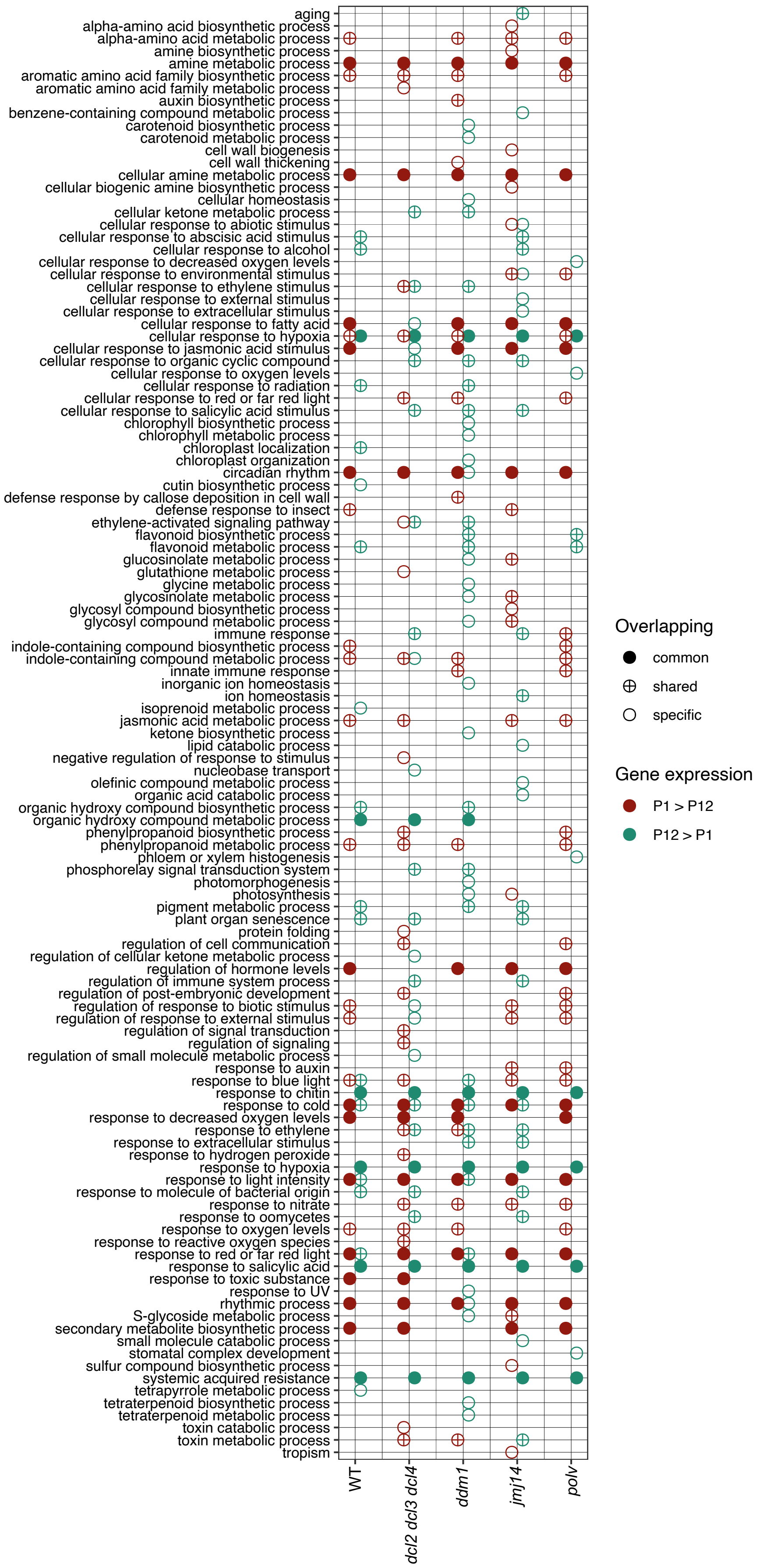


Supplementary Fig. S2

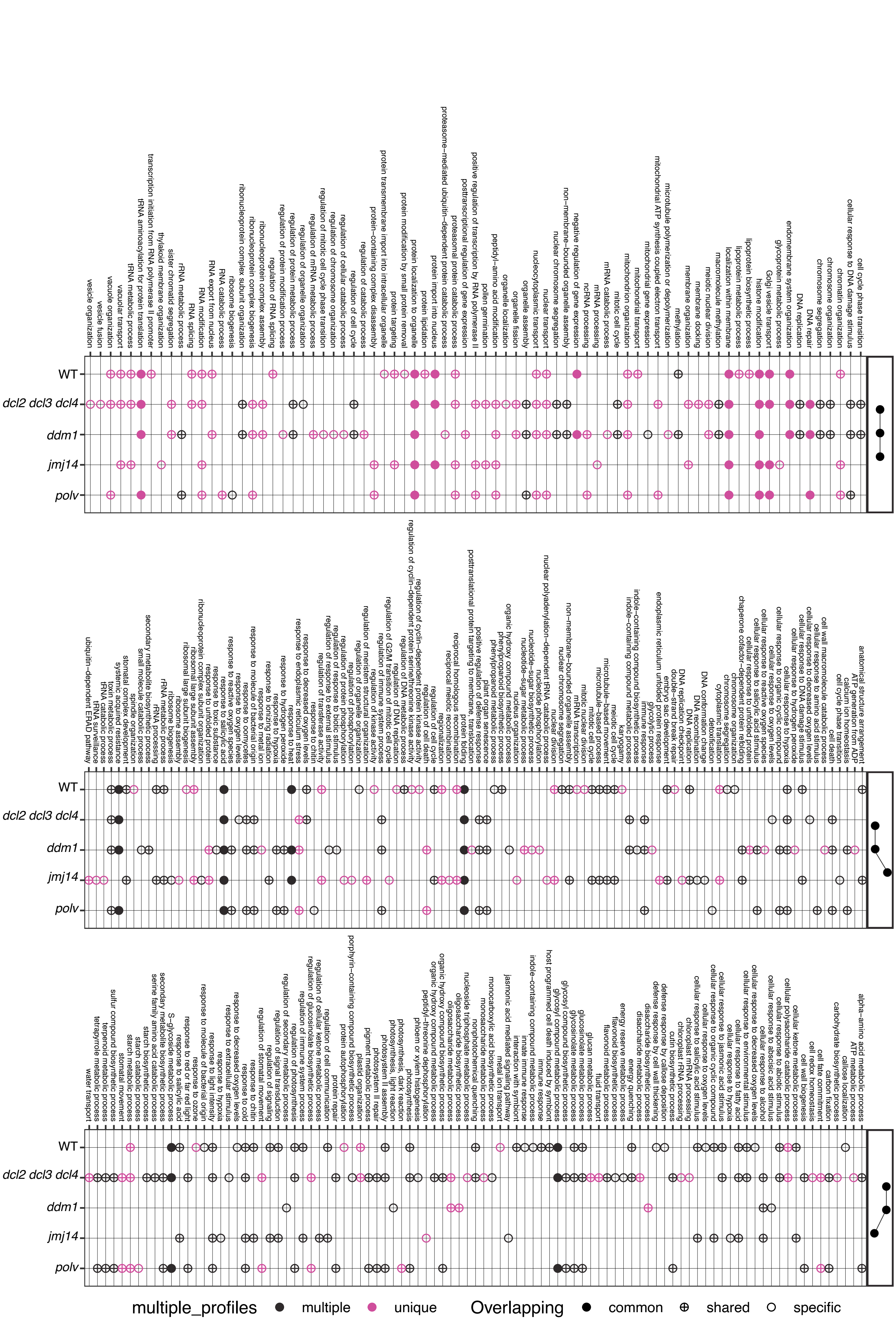


Supplementary Fig. S3

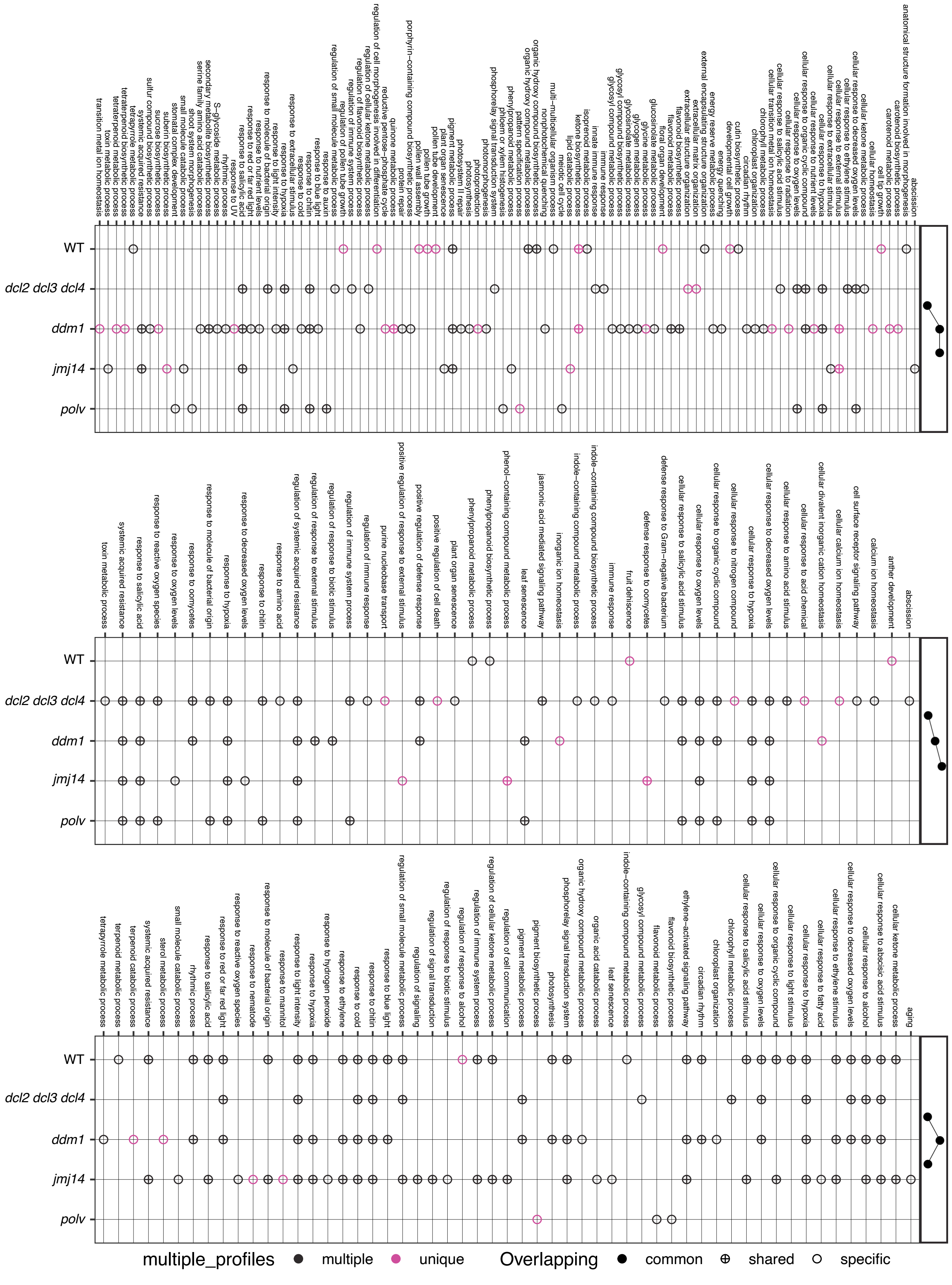


Supplementary Fig. S3 (cont.)

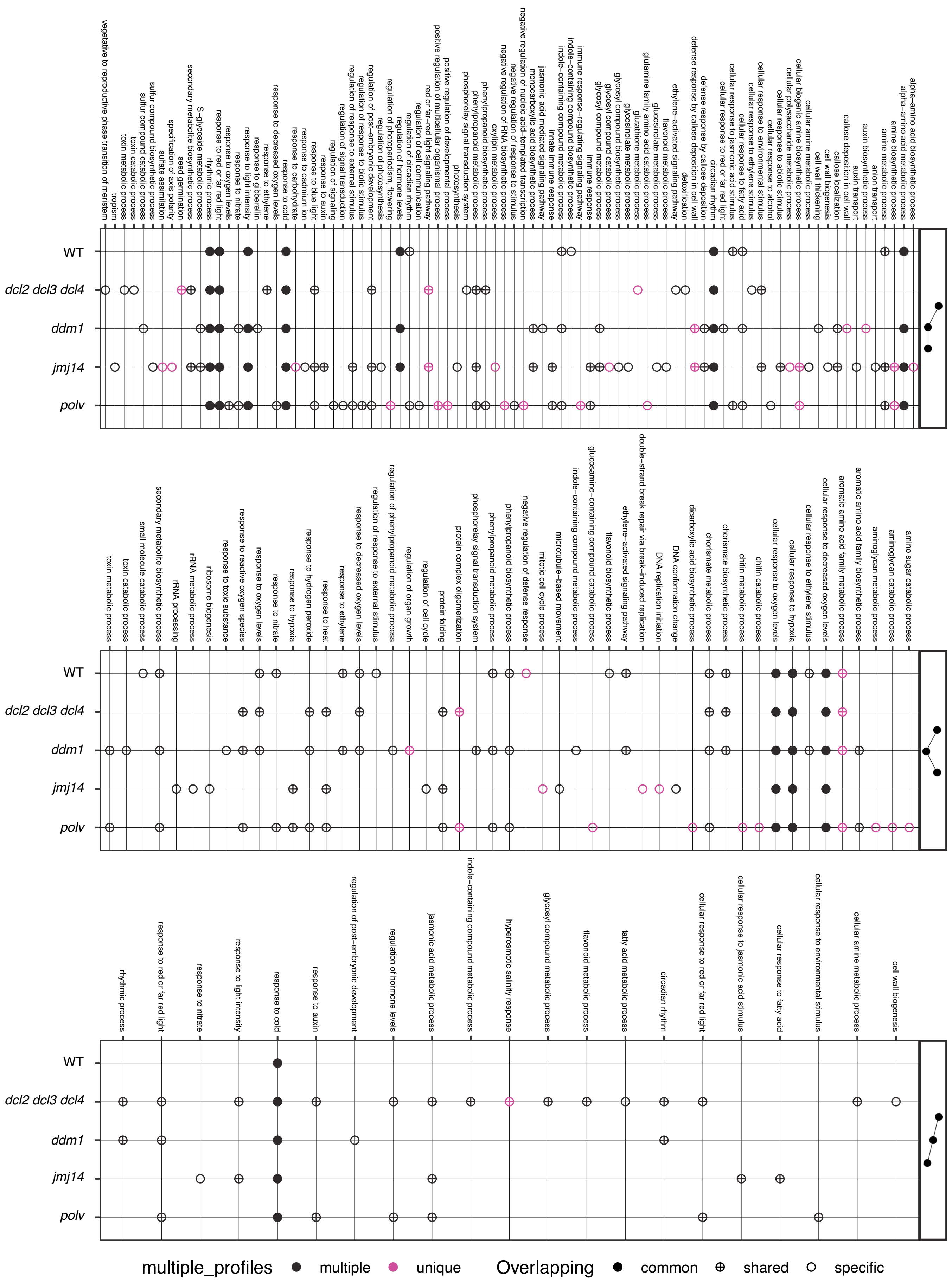


Supplementary Fig. S3 (cont.)

### Shared genes per profiles and genotype

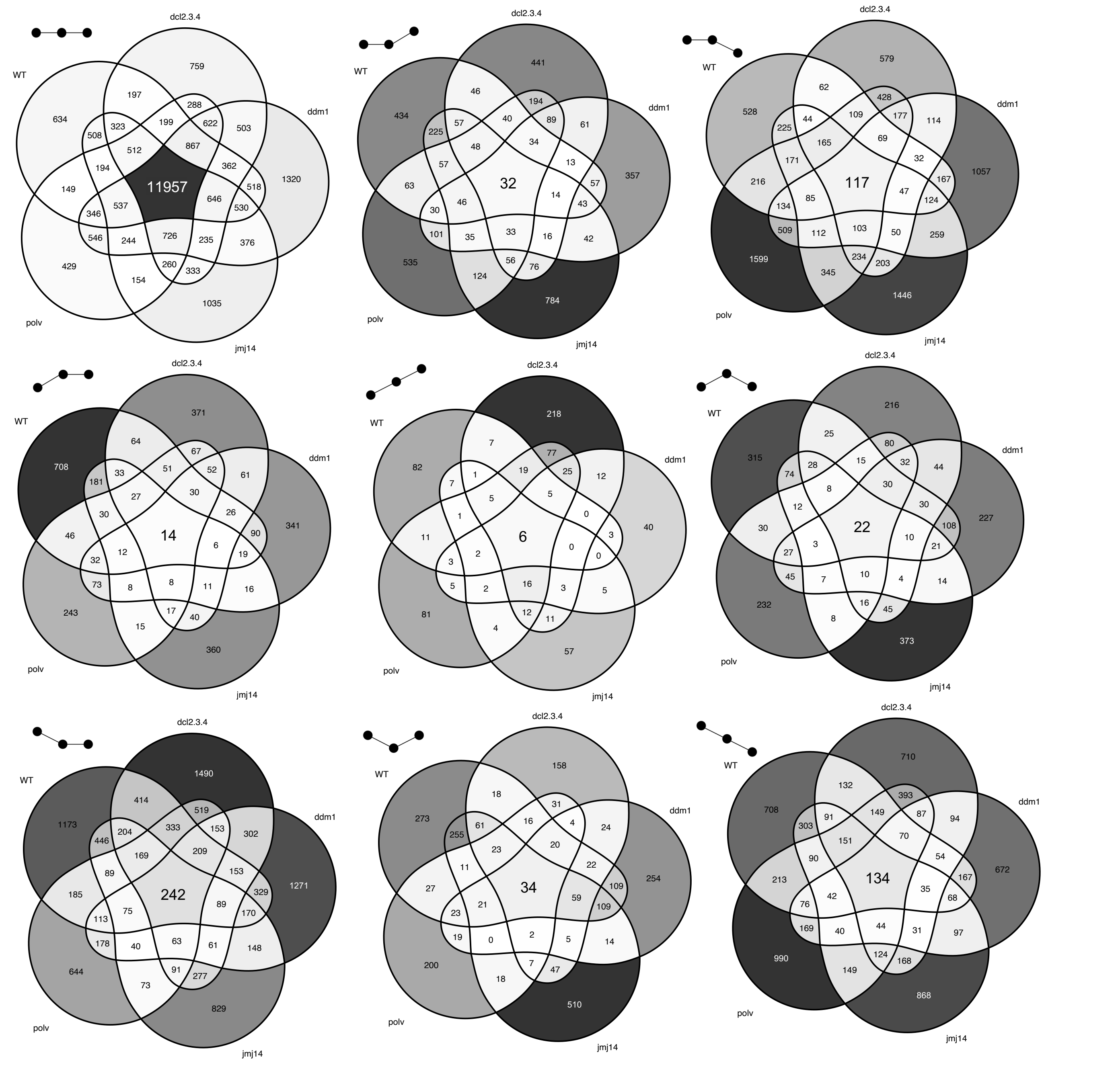


Supplementary Fig. S4

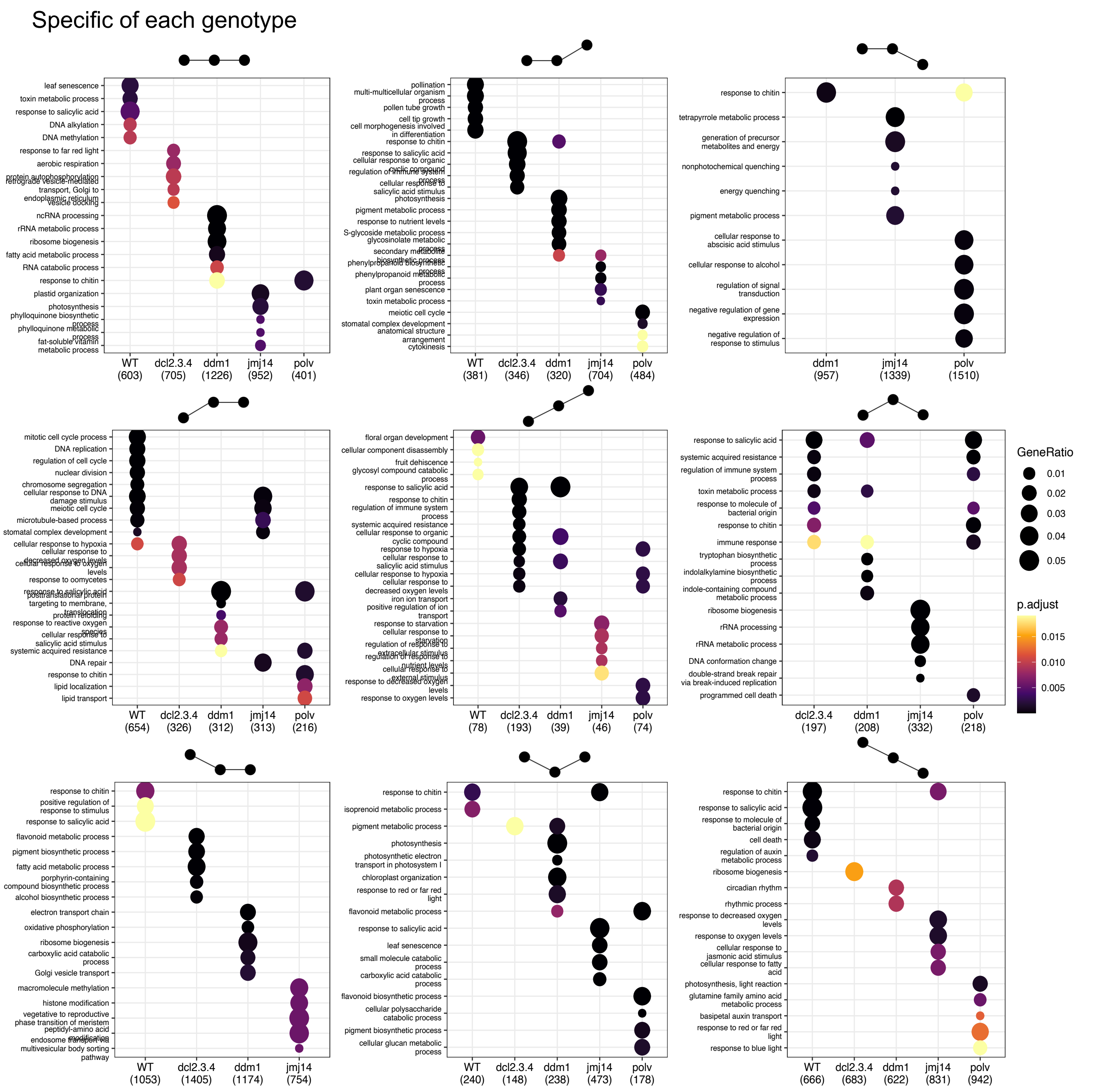


Supplemenary Fig. S5

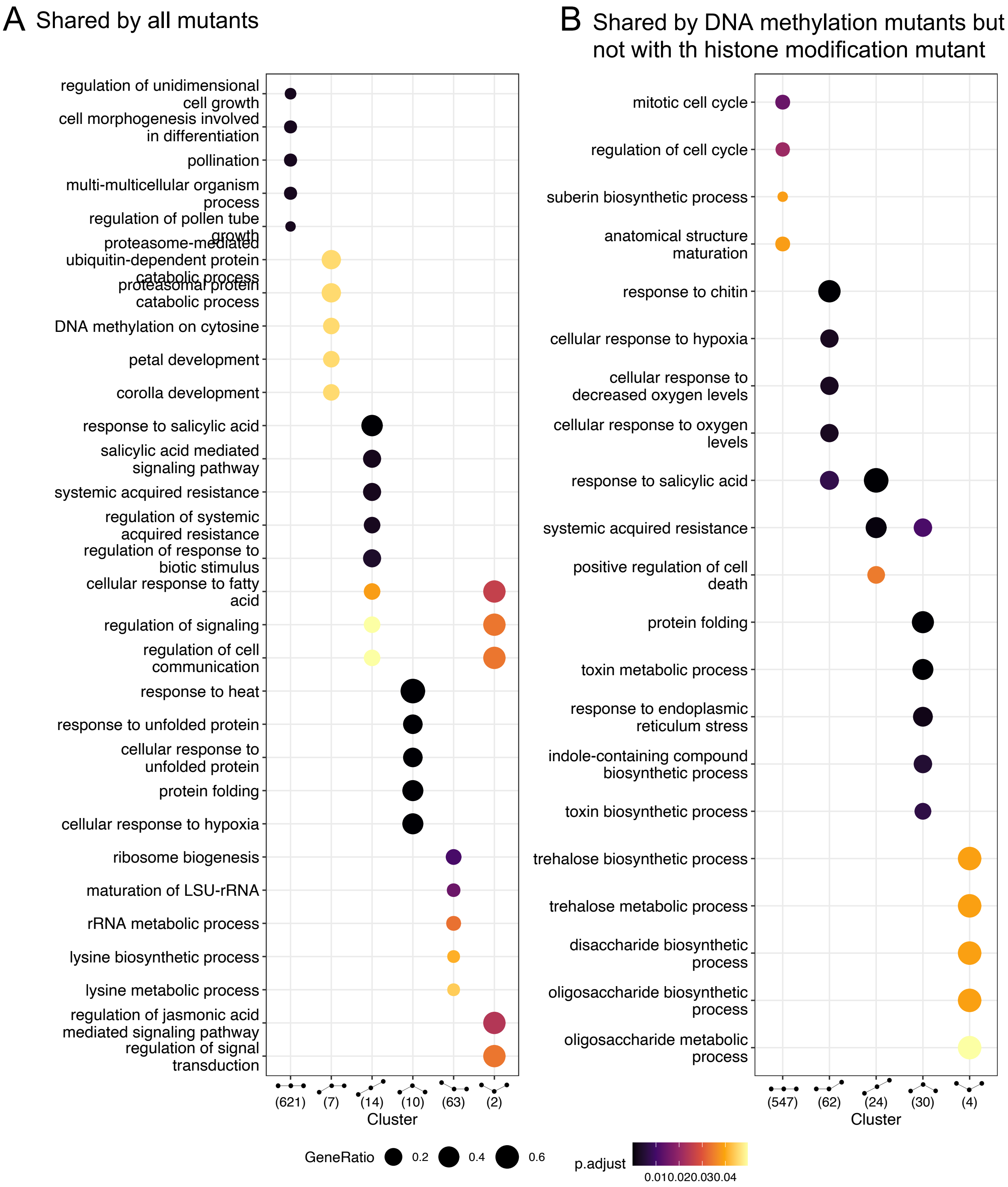


Supplementary Fig. S6

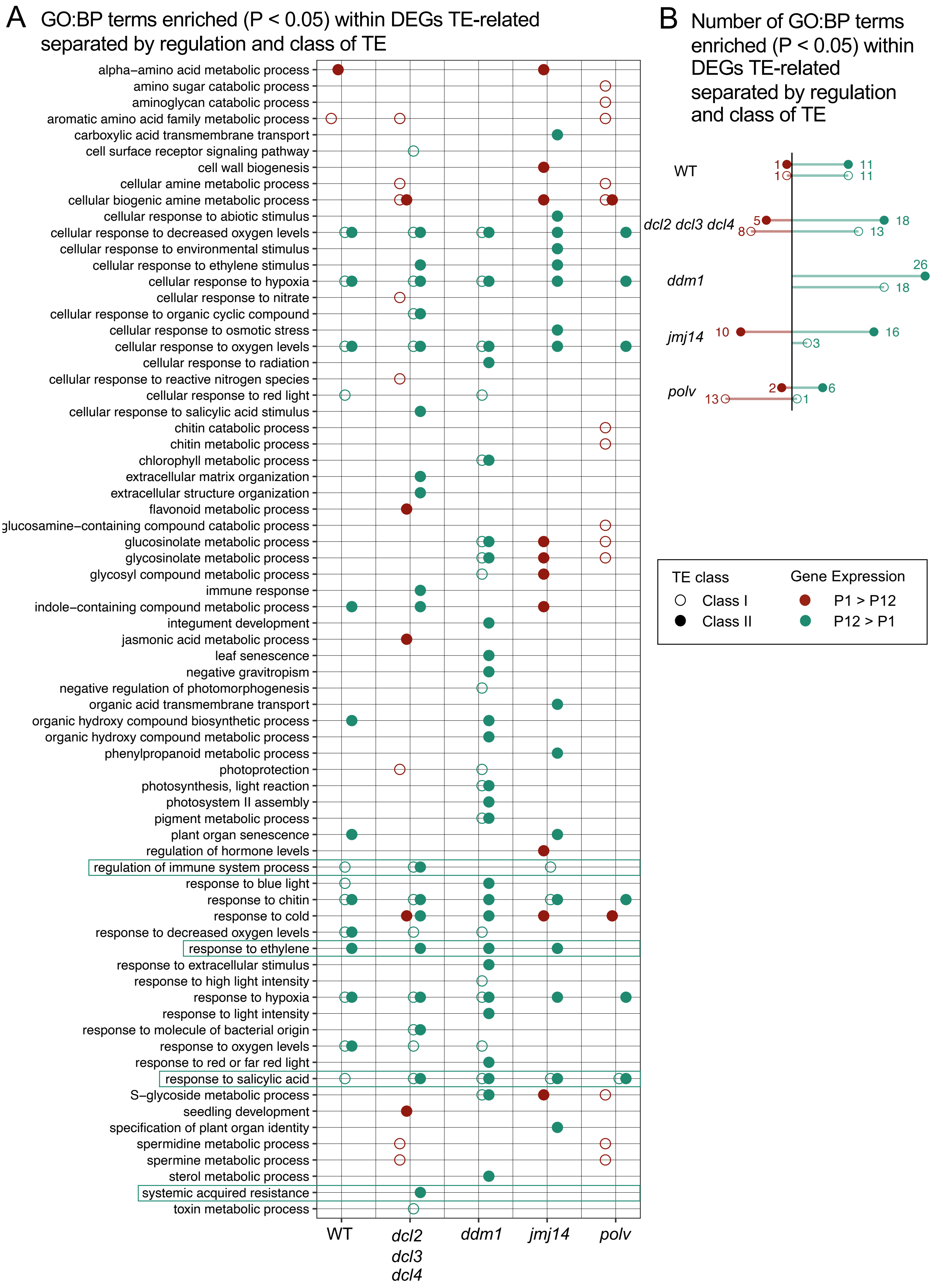


Supplementary Fig. S7

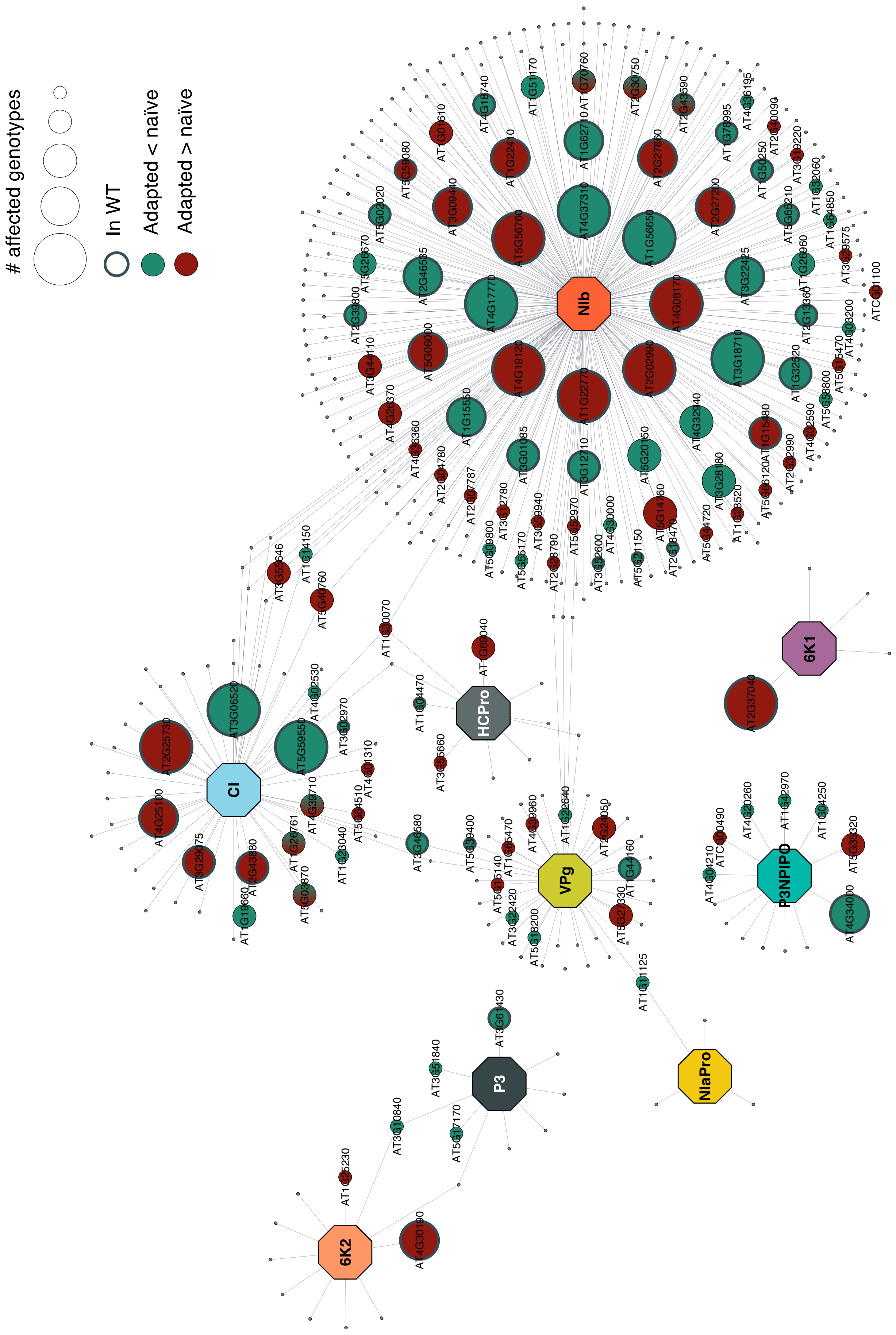


Supplementary Fig. S8

**Supplementary Table S1.** Genes with a differential gene expression in all the mutants respect WT under the same inoculation conditions.

| Conditions intersection | Gene (common name) | Description/Comments |
|---|---|---|
| **P12 ∩ P1 ∩ Mock** | AT1G53480 (*T3F20.20*) | *METHIONINE OVERACCUMULATION 1 (MTO1) RESPONDING DOWN 1* (MRD1) gene, and encodes for a protein involved in down-regulation of methionine biosynthesis, and is essential for salicylic acid-mediated defense (Blevins et al. 2014 ) |
| | AT2G01422 | Noncoding RNA unknown function. found in experiments seeking for candidate genes directly downregulated by RdDM-related siRNAs (Kurihara et al. 2008) |
| | AT3G28899 | Encodes for a member of the ribosomal protein L34e superfamily protein that has was already found in experiments seeking for candidate genes directly downregulated by RdDM-related siRNAs (Kurihara et al. 2008) |
| | AT4G04221 | Antisense long-noncoding RNA. negatively regulates the expression of *LEUCINE RICH-REPEART (LRR) RECEPTOR-LIKE PROTEIN 46* (*RLP46*), a plasma-membrane located receptor involved in tissue differentiation, plant development and senescence that has been shown to be upregulated after infection (Steidele and Stam 2021) |
| | AT4G04223 | Noncoding primary transcript of unknown function |
| | AT5G24240 (*PI4KC3*) | Ubiquitin family protein involved in regulation of flower development and responses to abscisic acid (Akhter et al. 2015) |
| **P12 ∩ P1 ∩ Mock** | AT1G22990 (*HIPP22*) | *HEAVY METAL ASSOCIATED ISOPRENYLATED PLANT PROTEIN 2*. HIPP metallochaperone proteins are involved in metal homeostasis and detoxification but also in transcriptional responses to temperature and drought and plant-fungi interactions (De Abreu-Neto et al. 2013) |
| | AT1G53490 (*HEI10*) | *ENHANCER OF CELL INVASION 10*. RING finger-containing protein of the ZMM class required for class I crossover formation during meiosis (Serra et al. 2018) |
| | AT3G09450 | Encodes for a member of the fusaric acid resistance family protein. This family of proteins is involved in detoxification of fusaric acid, a secondary metabolite produced during fungal infections (Bouizgarne et al. 2005). |
| | AT4G14140 (*MET2*) | *DNA METHYLTRANSFERASE 2*. Essential component of the DNA methylation that promotes gene silencing in response to genome defense against transposons and DNA viruses. Geminiviruses' Rep protein, for example, actively repress the expression of DNA methyltransferases *MET1* and *CHROMOMETHYLASE 3* (*CMT3*) in locally and systemically infected tissues (Rodríguez-Negrete et al. 2013). |
| | AT4G15390 (*DLI3740c*) | Encodes for a HXXXD-type BAHD acyltransferase family protein. BAHD acyltransferases are often involved in the production of phenolic secondary metabolites, and particularly in brassinosteroids homeostasis (Zhang and Xu 2018). Brassinosteroids play key roles in plant adaptation to infections (Xiong et al. 2022). |
| **Mock ∩ P1 − P1** | AT1G75945 | Mitochondrial membrane-associated hypothetical proteins |
| | AT2G33175 | Mitochondrial membrane-associated hypothetical proteins |


Akhter, S., Uddin, M.N., Jeong, I.S., Kim, D.W., Liu, X.M., & Bahk, J.D. (2015) Role of AtPI4Kgamma3, a type II phosphoinositide 4-kinase, in abiotic stress responses and floral transition. *Plant Biotechnol. J.*, **14**, 215-230.

Blevins, T., Pontvianne, F., Cocklin, R., Podicheti, R., Chandrasekhara, C., Yerneni, S., et al. (2014) A two-step process for epigenetic inheritance in Arabidopsis. *Mol. Cell.*, **54**, 30-42.

Bouizgarne, B., El-Maarouf-Bouteau, H., Frankart, C., Reboutier, D., Madiona, K., Pennarun, A.M., et al. (2005) Early physiological response of *Arabidopsis thaliana* cells to fusaric acid: toxic and signalling effects. *New Phytol.*, **169**, 209-218.

De Abreu-Neto, J.B., Turchetto-Zolet, A.C., Valter de Oliveira, L.F., Bodanese Zanettini, M.H., & Margis-Pinheiro, M. (2013) Heavy metal-associated isoprenylated plant protein (HIPP): characterization of a family of proteins exclusive to plants. *FEBS J.*, **280**, 1604-1616.

Kurihara, Y., Matsui, A., Kawashima, M., Kaminuma, E., Ishida, J., Morosawa, T., et al. (2008) Identification of the candidate genes regulated by RNA-directed DNA methylation in Arabidopsis. *Biochem. Biophys. Res. Commun.*, **376**, 553-557.

Rodríguez-Negrete, E., Lozano-Durán, R., Piedra-Aguilera, A., Cruzado, L., Bejarano, E.R., & Castillo, A.G. (2013) Geminivirus Rep protein interferes with the plant DNA methylation machinery and suppresses transcriptional gene silencing. *New Phytol.*, **199**, 464-475.

Serra, H., Lambing, C., Griffin, C.H., Topp, S.D., Nageswaran D.C., Underwood, C.J., et al. (2018) Massive crossover elevation via recombination of *HEI10* and *recq4a recq4b* during Arabidopsis meiosis. *Proc. Natl. Acad. Sci. USA*, **115**, 2437-2442.

Steidele, C.E., & Stam, R. (2021) Multi-omics approach highlights differences between RLP classes in *Arabidopsis thaliana. BMC Genomics*, **22**, 557.

Xiong, J., Wan, X., Ran, M., Xu, X., Chen, L., & Yang, F. (2022) Brassinosteroids positively regulate plant immunity via BRI1-EMS-SUPPRESSOR 1-mediated *GLUCAN SYNTHASE-LIKE 8* transcription. *Front. Plant Sci.*, **13**, 854899.

Zhang, Z., & Xu, L. (2018) Arabidopsis BRASSINOSTEROID INACTIVATOR 2 is a typical BAHD acyltransferase involved in brassinosteroid homeostasis. *J. Exp. Bot.*, **69**, 1925-1941.

**Supplementary Table S2.** Genes with opposed regulation within the core of DEGs involved in TuMV adaptation to *A. thaliana*

| Locus | Gene name | Regulation adapted virus (activated, repressed) | Description |
|---|---|---|---|
| AT1G69480 | *PHO1-H10* | WT (activated)<br>*dcl2 dcl3 dcl4* (repressed)<br>*ddm1* (activated)<br>*jmj14* (activated)<br>*polv* (activated) | Phosphate transporter PHO1 homolog 10; May transport inorganic phosphate (Pi); Belongs to the SYG1 (TC 2.A.94) family. |
| AT2G40610 | *EXPA8* | WT (activated)<br>*dcl2 dcl3 dcl4* (activated)<br>*ddm1* (activated)<br>*jmj14* (repressed)<br>*polv* (activated) | Expansin-A8; Causes loosening and extension of plant cell walls by disrupting non-covalent bonding between cellulose microfibrils and matrix glucans. No enzymatic activity has been found (By similarity). Belongs to the expansin family. Expansin A subfamily. |
| AT4G09820 | *TT8* | WT (activated)<br>*dcl2 dcl3 dcl4* (repressed)<br>*ddm1* (activated)<br>*jmj14* (activated)<br>*polv* (activated) | Transcription factor TT8; Transcription activator, when associated with MYB75/PAP1 or MYB90/PAP2. Involved in the control of flavonoid pigmentation. Plays a key role in regulating leucoanthocyanidin reductase (BANYULS) and dihydroflavonol-4-reductase (DFR). Not required for leucoanthocyanidin dioxygenase (LDOX) expression. |
| AT4G25480 | *DREB1A* | WT (activated)<br>*dcl2 dcl3 dcl4* (repressed)<br>*ddm1* (activated)<br>*jmj14* (repressed)<br>*polv* (repressed) | Dehydration-responsive element-binding protein 1A; Transcriptional activator that binds specifically to the DNA sequence 5'-[AG]CCGAC-3'. Binding to the C-repeat/DRE element mediates cold-inducible transcription. CBF/DREB1 factors play a key role in freezing tolerance and cold acclimation; Belongs to the AP2/ERF transcription factor family. ERF subfamily. |
| AT5G65730 | *XTH6* | WT (activated)<br>*dcl2 dcl3 dcl4* (activated)<br>*ddm1* (activated)<br>*jmj14* (repressed)<br>*polv* (activated) | Probable xyloglucan endotransglucosylase/hydrolase protein 6; Catalyzes xyloglucan endohydrolysis (XEH) and/or endotransglycosylation (XET). Cleaves and religates xyloglucan polymers, an essential constituent of the primary cell wall, and thereby participates in cell wall construction of growing tissues (By similarity); Belongs to the glycosyl hydrolase 16 family. XTH group 1 subfamily. |

**Supplementary Table S3.** Genes of the PPI network affected by viral adaptation in all plant genotypes.

| Viral protein | Plant protein | Common name | Description |
|---|---|---|---|
| **6K1** | AT2G37040 | PAL1 | Phenylalanine ammonia-lyase 1; This is a key enzyme of plant metabolism catalyzing the first reaction in the biosynthesis from L-phenylalanine of a wide variety of natural products based on the phenylpropane skeleton; Belongs to the PAL/histidase family. |
| **CI** | AT2G25730 | F3N11.1 | Zinc finger FYVE domain protein. |
| | AT3G06520 | F5E6.15 | Agenet domain-containing protein. |
| | AT5G59550 | DURF2 | E3 ubiquitin-protein ligase RDUF2; E3 ubiquitin-protein ligase involved in the positive regulation of abscisic acid-dependent drought stress responses. Possesses E3 ubiquitin ligase activity in vitro. |
| **NIb** | AT1G22770 | GI | Protein GIGANTEA; Involved in regulation of circadian rhythm and photoperiodic flowering. May play a role in maintenance of circadian amplitude and period length. Is involved in phytochrome B signaling. Stabilizes ADO3 and the circadian photoreceptor ADO1/ZTL. Regulates 'CONSTANS' (CO) in the long-day flowering pathway by modulating the ADO3-dependent protein stability of CDF1 and CDF2, but is not essential to activate CO transcription. Regulates, via the microRNA miR172, a CO-independent pathway that promotes photoperiodic flowering by inducing 'FLOWERING LOCUS T'. |
| | AT1G56650 | MYB75 | Transcription factor MYB75; Transcription activator, when associated with BHLH12/MYC1, EGL3, or GL3. Promotes the synthesis of phenylpropanoid-derived compounds such as anthocyanins and proanthocyanidin, probably together with GL3 and BHLH2. Regulates the expression of CHS, DFRA, LDOX, and BAN. |
| | AT2G02990 | RNS1 | Ribonuclease 1; May remobilize phosphate, particularly when cells senesce or when phosphate becomes limiting. |
| | AT3G18710 | PUB29 | U-box domain-containing protein 29; Functions as an E3 ubiquitin ligase. |
| | AT4G08170 | ITPK3 | Inositol-tetrakisphosphate 1-kinase 3; Kinase that can phosphorylate various inositol polyphosphate such as Ins(3,4,5,6)P4 or Ins(1,3,4)P3. Phosphorylates Ins(3,4,5,6)P4 to form InsP5. This reaction is thought to have regulatory importance, since Ins(3,4,5,6)P4 is an inhibitor of plasma membrane $Ca^{(2+)}$-activated $Cl^{(-)}$ channels, while Ins(1,3,4,5,6)P5 is not (By similarity). Also phosphorylates Ins(1,3,4)P3 or a racemic mixture of Ins(1,4,6)P3 and Ins(3,4,6)P3 to form InsP4. Ins(1,3,4,6)P4 is an essential molecule in the hexakisphosphate (InsP6) pathway (By similarity). |
| | AT4G17770 | TPS5 | Alpha,alpha-trehalose-phosphate synthase [UDP-forming] 5; In the N-terminal section; belongs to the glycosyltransferase 20 family. |
| | AT4G19120 | ERD3 | Probable methyltransferase PMT21; Belongs to the methyltransferase superfamily. |
| | AT4G37310 | C7A10.50 | Cytochrome P450, family 81, subfamily H, polypeptide 1; Belongs to the cytochrome P450 family. |
| | AT5G56760 | SAT5 | Serine acetyltransferase 5; Belongs to the transferase hexapeptide repeat family. |
